# A Darwinian model of cultural replication over the continuum of Weismannian and Lamarckian inheritance

**DOI:** 10.1101/2024.10.09.617348

**Authors:** István Zachar, Szabolcs Számadó

## Abstract

Universal Darwinism argues that entities capable of multiplication, variation and inheritance evolve by natural selection. Besides genetic reproduction implementing Weismannian inheritance, other adaptive systems were proposed to support Darwinian evolution, most notably culture, assuming autonomous cultural replicators akin to genes. Culture, on one hand, lacks direct template copying and inherits acquired changes, while on the other its overall behaviour is Darwinian as languages attest. The properties of culture setting it apart from biological evolution suggests a Lamarckian, while its overall behaviour a Weismannian interpretation, which is often coined as “the duality of cultural replication”. According to canonical Darwinism, Lamarckian and Weismannian inheritance are disjunct mechanisms and only the latter is sufficient to maintain cumulative evolution. We resolve this apparent contradiction via a mathematical model of cultural information replication. We model the inheritance of information, cultural or genetic, as a successive set of template-based transformations of digital information between generations. We demonstrate that Lamarckian and Weismannian inheritance form a continuum and that both can reconstruct information stably for an indefinite time. Our formalism naturally unifies Lamarckian and Weismannian inheritance, proving that cultural replication, in theory, can approximate the fidelity of genetic replication. We claim that (well-defined) Lamarckian inheritance is an adequate low-level model of culture and can support population-level Darwinian evolution. We discuss the implications of our results to bridge the gap between the Lamarckian and Darwinian explanations of cultural evolution.

**Significance statement:** Much of the standstill of cultural evolution theory stems from under-defined concepts that lack rigorous mathematical formalism. We provide a strict application of established concepts of biological evolution to the replication of cultural information. According to our analogy, cultural replication corresponds to Lamarckian inheritance (inheriting acquired changes), genetic replication to Weismannian inheritance, both being part of the same continuum of informational replication. We prove in our model, that despite a lack of template copying in Lamarckian replication, overall copying fidelity can approximate that of Weismannian replication, i.e. culture can approximate genetic copying fidelity. Cultural inheritance may be Lamarckian at the level of inheritance (microevolution) while it can still lead to Darwinian evolution at the population level (macroevolution).

## Introduction

Biological organisms are units of evolution that not only reproduce from time to time but also evolve by inheriting variations. The theory of Darwinian evolution posits that entities able to multiply, acquire variations and inherit these variations will evolve by means of natural selection ^1–6^. The mechanism of genetic inheritance behind the evolution of organisms is accounted for by well understood molecular mechanisms. However, as recognized by Universal Darwinism ^7–10^, the above conditions of evolution are general enough to be applicable to any complex adaptive system, not exclusive to molecules, genes, and organisms. Therefore, Darwinian evolution is not restricted to biological organisms and arises whenever the three conditions are met ^8^. We now understand that Darwinian dynamics is responsible for evolution in many complex adaptive systems ^11,12^: cellular inheritance systems ^13,14^, the adaptive immune system ^15,16^, niche inheritance ^17,18^, replicative computer programs ^19^ potentially neuronal patterns or even cognitive functions in human brains ^20–22^, animal culture ^23–25^, and most notably, human culture ^26–32^ (for a review, see ^33^) and language/grammar ^34– 37.^

Dawkins drew an analogy between genetic and cultural inheritance, from the replicator’s eye view according to which cultural evolution is shaped by autonomous cultural replicators (‘memes’) akin to genes governing organismal evolution ^27,38–41^. At that time, the establishment of memetics as a validated theory (practically as Modern Synthesis applied to culture) seemed within reach ^38,42,43^. However, the peculiarities of culture, and the marked differences compared to genetic inheritance, hindered progress toward a formal model on a par with genetics.

Cultural inheritance, while obviously relying on genes determining cognitive capabilities, is considered largely independent of genetic inheritance, forming thus one leg of a *dual inheritance system* ^31,44–46^ (also see ^47,48^). During (social) learning, information is inherited in an entirely different way of the clear molecular specificities of genetics, without a well-characterized underlying copying mechanism (template replication, i.e. the bitwise copying of nucleotides from a template to a new copy) and without a single informational parent. Culture is not only acquired from cultural parents but can also be obtained from cultural artifacts. In general, cultural information is inherited from potentially many sources, including conspecifics or objects (see Figure 1). As J.B.S Haldane has aptly noted, the word ‘inherit’ is quite unfortunate to denote property-acquisition from a parent, as it can mean both a character inherited genetically (baldness) but also an object acquired “legally” (a watch) ^49^. This dichotomy, characteristic of culture, lead to a long-standing debate on *what* exactly is replicated by culture ^20,26–28,30,39,50–56^ and whether Darwinism (specifically, memetics) can serve as an analogy for cultural evolution at all ^29,47,57–60^. Attempts to provide an explanation of cultural phenomena via Darwinian mechanisms were challenged, and the progress of memetics has stalled ^57,61,62^. However, insisting on a discrete unit of cultural evolution comparable to a gene may have been a misdirection considerably contributing to the failure of memetics ^63^.

**Figure 1.**
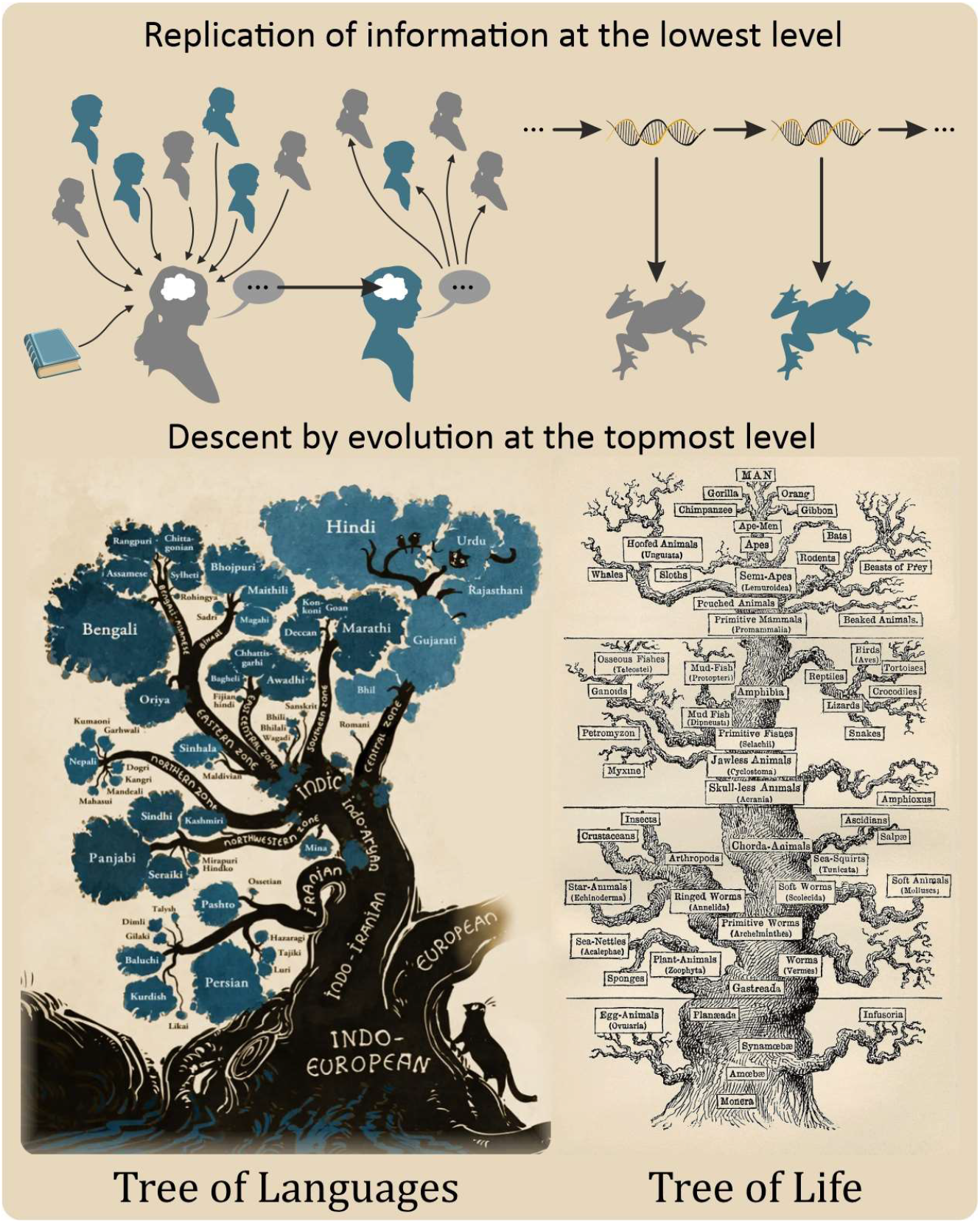
The duality of cultural inheritance. The replication of cultural information at the lowest level of individual interactions (top row) does not follow the strict genetic replication of information of biological organisms. Nevertheless, the result of an indefinitely long process resembles a tree, similar to the tree of organisms generated by biological evolution, when observed from the surface level (bottom row). Tree of Languages image is courtesy of Minna Sundberg, Tree of Life is by Ernst Haeckel, from Wikipedia public domain, originally from ^67^.

The situation has not improved since regarding valid candidates of cultural replicators or models of cultural evolution that can effectively bridge the gap to their corresponding biological (Darwinian) concepts. While Brewer et al. have identified “modelling culture as a complex adaptive system” as a major challenge of cultural evolution theory ^64^, progress is hindered because cultural evolution research (theory, experimentation and modelling alike) often happen without a strict, formal definition of concepts involved ^65^. As cultural evolution often uses the same vocabulary as more conventional biological evolution, the lack of formal clarity consequently and inevitably leads to confusion and standstill (see ^66^).

Much of the difficulty follows from the purported *Lamarckian nature of cultural evolution* ^38^. To understand the difference between the Lamarckian and Darwinian approaches to cultural inheritance, one must understand the nature of cultural information reproduction. Cloak has distinguished *internal* from *material culture*, pointing out that internal culture consists of “instructions”, encoded in neuronal material, that do not get copied directly ^68^ (see Figure 2C). Because material culture (behaviour, rituals, social organization, tools, technology, etc.) is external to the human brain and includes non-material items too (e.g., gestures, utterances), it seems more appropriate to refer to it as *external*, being the antonym of internal. Human culture is a mixture of both internal and external items, that together are responsible for the transmission and maintenance of cultural information. In this sense, culture clearly differs from biological evolution, where, according to the Central Dogma of biology, the genetic information is maintained exclusively by the successive copying of genes without inheriting information from the body ^69^ (Figure 2A,B). There are many exceptions (e.g. epigenetic inheritance systems ^70–72^), but in principle, it ensures the stable replication of information and reproduction of organisms.

**Figure 2.**
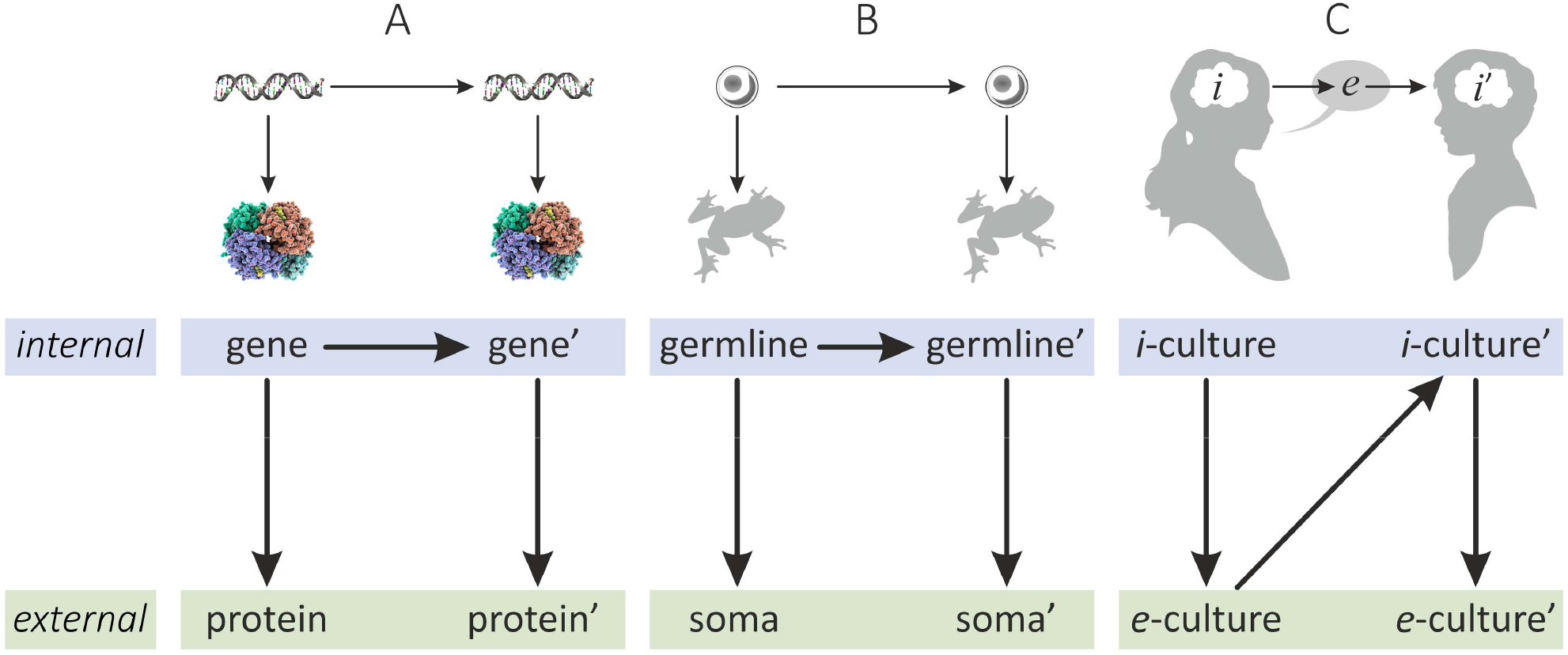
Simplified information inheritance topologies. Arrows indicate the flow of information. **A**: **Genetic inheritance**. Genes are copied directly but seldom interact with the selective environment directly. Proteins, translated from DNA, are under functional constraints and selection. **B**: **Organismal inheritance**. Inheritance is Weismannian as cells of the germline replicate without being under direct selection. A single germline cell (the fertilized egg) defines the body (soma) of the organism. in more general terms, the phenotypic properties of the organism are defined by the genome, the sum of all heritable traits and only the genome can inherit acquired changes. The process rests on the copying of the genome encoded in the DNA of the germline to yield a potentially different genome (germline’). There is a division of labour between the two states, often termed replicator and interactor ^27,73^. The lineage of replicators (blue) is hidden from direct external effects of the environment; selection directly affect interactors (green). Selection only indirectly affects the survival and distribution of replicators. **C**: **Cultural inheritance**. En element of internal (i-)culture is inherited via intermediate external (e-)cultureelement to yield a new element of i-culture. For sake of simplicity, we call the host of i the teacher and the target of the transmission the learner.

The lack of a similar principle in culture led to the claim that, unlike the obviously Darwinian evolution of genes, cultural evolution is inherently Lamarckian ^13,26,38,74^. However, ‘Lamarckian’ is an ill-defined concept, meaning different things at different authors ^13,75–77^. It could imply the inheritance of acquired characters ^13^, directed evolution by use and disuse ^78^, or an inherent drive for complexity ^76,79^. Some have argued that cultural evolution need not be Lamarckian as it does not rely on the genetic inheritance system but social transformation (cf. ^76,80,81^) and that Lamarckism would explicitly require a dedicated genotype and phenotype ^79^. Applying the terminology of genetic inheritance to culture without a proper mapping meant that talks about cultural “evolution” remained a superficial analogy to biological evolution.

Nevertheless, epigenetic inheritance, culture included, is often defined as having a ‘Lamarckian flavour’ as some changes during the lifetime of the individual can be inherited, yet such inheritance is not governed by template copying of replicators ^13,72,82^. For the modelling purpose of this paper, we define Lamarckian inheritance as an information transmission mechanism that lacks template copying and can potentially inherit acquired changes of the soma (state under selection). On one hand, culture lacks a direct copying mechanism (from brain to brain) or a dedicated singular parent and replication event. It seems to entail that, compared to DNA replication, cultural transmission has low fidelity, especially in animals ^83^, and that information is inevitably lost in the process. Consequently, it may seem that cultural transmission cannot support Darwinian evolution. On the other hand, culture clearly has all the hallmarks of complex adaptive Darwinian systems: reproduction, variation, and inheritance ^84^ and cultural patterns are often stably maintained, as religions and languages attest, which requires the continued, stable replication of information ^85–87^. This apparent contradiction is called the **duality of cultural inheritance**, the phenomenon that culture seems to be Lamarckian locally at the micro-scale of individual transmission events (“the information encoded is […] not readily dissociated into independently heritable parts, but is contained in the dynamics of the interactions between the organism and its social and ecological environment” ^13^), yet Darwinian at macro-scale ^84,88^ (see Figure 1). Thus, while genetic inheritance is overwhelmingly Weismannian to stably maintain information, Lamarckian inheritance is considered insufficient to maintain cumulative evolution, leading to an apparent contradiction regarding how culture can evolve and maintain information stably.

There are not many models that aimed to capture social learning and cultural inheritance in the evolutionary context. Cavalli-Sforza and Feldman, through a series of gene–culture coevolutionary models ^44,89–94^ converged to a simple representation of cultural inheritance examining the contribution of cultural transmission to genetic traits ^92^. Why their pioneering work was influential to establish the notion that cultural traits can be inherited stably, parallel to and phenomenologically similar to genetic traits (or even dominating over them), we believe, that what Cavalli-Sforza and Feldman have investigated was only one pillar: the *population genetics of cultural inheritance*. We took a deeper look at the complicated processes behind simple transmission rates of trait types, by investigating a more elementary aspect of cultural information transmission: the *information theory of cultural inheritance (inheritance topology)*. We are interested in how digital, compositional information is quantified, inherited and maintained through a culture-specific inheritance topology. We have ignored population-level factors (teacher-learner structure, selection, gene-culture effects, etc.) and have assumed that the transmission (replication, change) of one cultural trait to another is defined by multiple underlying informational states and their couplings. Practically, we have approximated cultural inheritance to digital template-copying-like processes, but with a different informational topology compared to genetic inheritance (Figure 2).

Culture is often considered an additional medium of evolution besides genetic. In particular, language seems to be a prime example of a complex adaptive Darwinian system that can maintain cultural evolution ^34–36,95^. It has a digital and compositional structure, an imitation-based learning mechanism clearly approximating copying, emergent linguistic variations and an apparent spreading and fixation of these variations in linguistic populations (for a description of how language learning compares to replication, see Figure SI 12 and Appendix 7). However, culture exists not only in humans but in animals too, albeit lacking a complex, symbolic language (e.g. primates ^96–98^, cetaceans ^99–101^, birds ^23,102^, and others ^103^). Therefore, a general explanation of cultural replication must be independent of human language and its digital, symbolic medium.

It is well known, that Bayes’ theorem (or Bayesian update, inference) is formally equivalent to a mathematical description of natural selection at the phenotypes’ level, e.g. via Price’s equation ^104–106^, as the Price equation is equivalent to the replicator equation of discrete phenotypes ^107^. Bayesian updatehas been applied to various contexts to explain complex adaptive behavior, e.g. to multiple levels of selection and structure learning ^108^, and notably, to culture ^7,109,110^. Thus, Bayesian update (or any of its equivalents) can describe natural selection of cultural information. It naturally follows then, that the same formal model of informational replication applicable to genetic inheritance should be applicable to cultural inheritance due to the top-level analogy, despite low-level discrepancies. We therefore applied the information- (template-) replication framework to cultural replication.

We argue that the main reason behind the duality of cultural inheritance, and the subsequent debate about a possible Darwinian account of culture, is the lack of a unified model that can account for both biological (Weismannian) and cultural (Lamarckian) inheritance. According to the distinction of internal and external culture by ^68^ (also see ^111^, who differentiate between copyable observable behaviour and non-copyable mental culture), there is a clear analogy between the germline/soma distinction and the internal/external levels of culture, with a distinct topology (Figure 2). Here we provide a unifying model, formalizing this analogy, following previous work ^4^. We formally define the distinct inheritance modes: direct, Lamarckian and Weismannian inheritance. We show that they are part of the same topological continuum, and in theory, all three can yield stable replication and, consequently, Darwinian evolution, regardless of the particulars of underlying mechanisms and whether there is direct template copying or not. We describe the necessary conditions for information to be stably maintained by Lamarckian inheritance. We discuss implications for cultural, and especially for language evolution. Finally, we establish an equivalence between the biological and cultural domains to facilitate crosstalk across disciplines.

## Results

To clarify the distinction between Lamarckian and Darwinian inheritance methods, we rely on the concepts of differentiated germline and soma, as instances of the more abstract concepts of genotype and phenotype (Figure 2, also see Table SI 1 for a mapping of terms between biological and cultural evolution). The phenotype refers to the expressed features of an organism (body, height, colour, etc.), whereas the genotype refers to all genes hidden inside and inherited by new generations (Figure 2A). The genotype-phenotype distinction has appeared precursory in the work of August Weismann ^112^ who argued that the reproductive cells form a distinct lineage hidden from external effects (the *germline*) opposed to the lineage of vegetative cells (*soma*) that is subject to environmental effects and can thus acquire changes but practically cannot pass them on (Figure 2B). Weismannian inheritance is a natural extension of the genotype-phenotype distinction of an evolutionary trait to the germline-soma distinction at the organismal level. Weismann argued that there is no information flow from the soma to the germ line, therefore acquired characters are not inherited ^112^. However, when the concepts of germline/genotype and soma/phenotype are mapped to cultural transmission, it becomes evident that the *topology* of cultural inheritance differs qualitatively from that of Weismannian (Figure 2). While in theory, any cultural transmission could be approximated by a template-based transmission (i.e. a process where a template is read bit-by-bit to generate a new copy), cultural replication can inherit acquired changes of *both* states (internal or external), corresponding to Lamarckian inheritance. That is, there is *always* an intermediate external medium under direct selection, as Cloak has recognized it ^68^ (Figure 2C). The difference in fundamental inheritance topologies justifies the different treatment of replication processes in our model.

The analogy rests on the similar functions encompassed by genes and bodies vs. internal and external cultural items. In general, the genotype and phenotype relate in two ways: the genotype is directly responsible for the development of the phenotype, while the phenotype indirectly affects the survival and distribution of genotypes in descendant generations. The phenotype is a representation of the genotype in any selective environment, thus it is also subject to selection, which, indirectly, affects the selection and distribution of genes. This idea was formalized by David Hull who introduced the concept of the *interactor* as “an entity that interacts as a cohesive whole with its environment in such a way that this interaction causes replication to be differential” ^73^. In biology the genotype refers to the genetic information being replicated, corresponding to the evolutionary replicator, while the body (soma) is the manifestation of the phenotype ^4^. Either can maintain information, but only one is under direct selection (Figure 2 bottom row).

We model the inheritance of information as a digital transmission process. During transformations from one state to the other, information may be copied, compressed, reconstructed, and possibly lost ^113^, as being mapped between different domains (i.e. media with different inventory sets). The route of information along these transformations from state to state defines the informational topology of inheritance (Figures 2, 3). In our formalism, we only consider informational replicators ^114,115^, as non-informational or limited hereditary ^116^ replicators cannot inherit sufficient amount of information for evolution (biological or cultural) to be open ended or interesting ^4^. Therefore, we assume that, regardless of the general lack of template replication, cultural inheritance does maintain sufficient amount of information. This means that, regardless of low-level mechanisms, cultural inheritance can be formally approximated and characterized by a digital process where bits of information are transmitted between generations and media akin to how biosequences are copied, transcribed, and translated (Appendix 7 gives an example how language can be considered a digital inheritance system comparable to genetic replication). Therefore, we define a digital inheritance system as a set of distinct states, encoding information in distinct media, linked by elementary digital transformations, forming an informational topology. The amount of information that can be stably maintained and transmitted for an indefinite time (if any) defines the *hereditary potential* of the system ^116,117^. Since the inheritance of evolutionary replicators is about maintaining information, the flow along the informational topology is analysed from the viewpoint of information loss, and we ignore material or energetic topologies, which may not follow the same route as information (e.g., DNA replication is semiconservative, where parental material is shared between daughter lineages).

**Figure 3.**
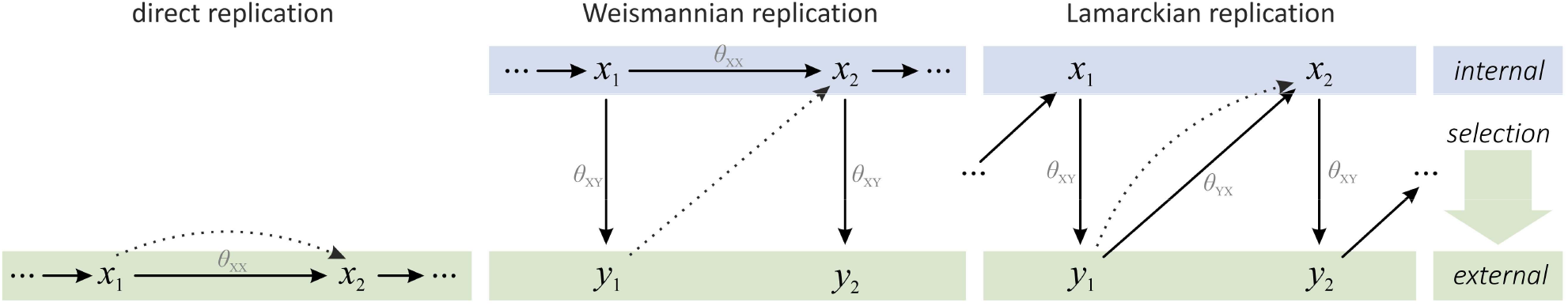
Distinct inheritance methods and their topologies of information transmission under selection. The x_i_ ∈X represent internal states that can presumably conserve information (genes or elements of internal culture, blue boxes) and are transmitted (inherited, copied or learnt) from individual to individual). The y_i_ ∈ Y represent external states of the replicator (soma or elements of external culture) that directly interface the environment and selection (green boxes). Normal arrows indicate transformation of information, dashed arrows indicate the indirect effect of selection on the distribution of the next generation of replicators. θ_XX_ is copying, θ_XY_ is translation, θ_YX_ is the reverse of translation. Note, that in case of direct replication, there is only one state that itself conserves information and is subject to direct selection, without a division of labour between different states, therefore it does not matter whether we call it x or y (cf. antidiagonal corners in Figure 6).

We have analysed all possible informational topologies of iterative transformations up to 2 distinct states (internal and external, X and Y), 4 elementary transformations (one-to-many, many-to-one, one-to-one and identity; see Appendix 2), assuming that selection acts directly on one state only and there is a single state contributing to the next generation (Figure 4). For details, see Methods and Appendix 1. In turn, we explain our conclusions drawn from this formal model.

**Figure 4.**
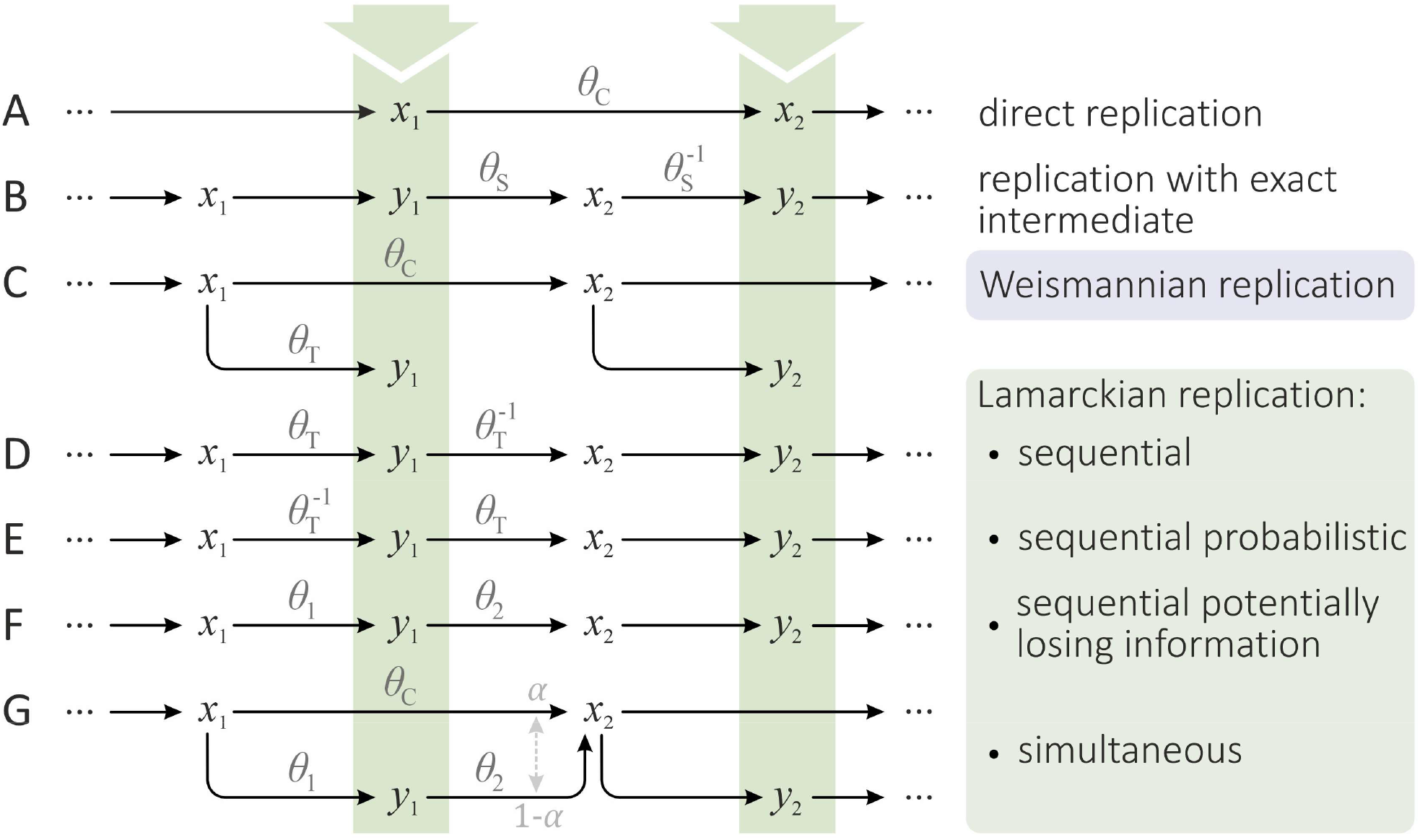
Possible specific topologies of information transmission over two distinct information-bearing states, x_i_ ∈ X and y_i_ ∈ Y, x denoteing successive internal and y denoting successive external states. Each letter stands for a new generation of the state; identical letters indicate structural identity of successive states. Selection (green arrows) acts on the green-shaded components. Stoichiometry is ignored, but one state is always assumed to be autocatalytic. Ignoring mutation and selection, information can be lost due to the reductive nature of elementary transformation functions, θ, which can be bijective (θ_C_, θ_s_), surjective but non-injective (θ_t_) or multivalued 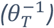 **A**: Without an Y state, when all successive θ are copying, the replication is trivial and direct (cf. Figure 3A). **B**: When successive θ-s map 1-to-1 and are inverses of each other, the intermediate y_t_ always exactly represents x_t_. **C**: In Weismannian replication, the Y state does not take part in inheritance at all (cf. Figure 3B). **D, E, F**: In sequential Lamarckian replication, y_t_ is defined solely by x_t_. Depending on the nature of successive transmissions θ, replication can be exact or inherently losing information (cf. Figure 3C). **G**: In the most general Lamarckian case, both x_t_ and y_t_ may simultaneously define x_t+1_under the mixing ratio of α. For more information, see Methods and Appendix 1.

### 1. Transmission states are all equivalent under selection

A crucially important condition of any such potentially multistate inheritance system is the **selective equivalence principle**, introduced in ^4^. According to the principle, to maintain evolutionary information for an indefinite time, the minimal condition is that successive states are equivalent under selection, that is *x*_l_ ∼ *y*_l_ ∼ *x*_2_ ∼ *y*_2_ ∼ …. where ∼ denotes selective equivalency. Equivalency means that phenotypes of successive states are indistinguishable under selection, regardless of their informational (or material, etc.) differences). For example, the gene *x*_l_ has a phenotype realized as *y*_l_, and – assuming no mutation, sexual recombination, epistasis, etc. – the gene *x*_2_ copied from *x*_l_ has an equivalent phenotype realized as *y*_2_.

Note, that while informational identity ensures phenotypic equivalence, successive states need not be informationally identical to maintain equivalency under selection. There may be neutral differences both among the elements of X or of Y, depending on the nature of transmissions X → X or X → Y. For example, neutral mutants may yield the same phenotype in a given environment. One consequence is that successive states may not store the same amount of *potential heritable information*, as their mutational distribution could differ significantly. Hence, states are not necessarily equivalent under evolution. Continuous evolutionary equivalence of replicators would exclude potential for open ended evolution.

### 2. There are only three qualitatively different replication topologies

An important claim of ours is that these inheritance methods can only be defined in terms of informational topology, of division of labour (storing information and interfacing the environment), and of information loss. That is, direct, Weismannian and Lamarckian inheritance are only defined for *informational replicators* ^114,115^, replicators that can be represented digitally (though need not be in reality). Consequently, the respective evolutionary units are direct, Weismannian or Lamarckian replicators.

A crucial observation is, that the division of labour of functionalities between *x* ∈ X and *y* ∈ Y is not necessarily exclusive. In case of Lamarckian replication (e.g. in Figure 3C), where *x* and *y* states alternate successively during inheritance, state *y* is the subject of direct selection but **both *x* and *y* pass on acquired changes** as heritable information. One can, however, imagine topologies where functionalities are further shared between entities. Some theories of cultural inheritance assume, that in some cases the external state may directly contribute to the next generation of external states (e.g. ‘contagion’, see ^79^). When the external state *y* inherits information, either 1) it does it exclusively or 2) it shares inheritance with the internal state *x*. In case 1), *y* is the *de facto* genotype and *x* has no role at all in selection or evolution and the system simplifies to direct replication *y*_l_ → *y*_2_ (Figure 3A). In case 2), both *x* and *y* contribute to the genotype and the topology simplifies again to direct replication, where a combination *f*(*x*_*t*_, *y*_*t*_) defines *x*_*t*+1_ (Figure 3G). We have specifically tested how such a system behaves under selection, applied on the Y state (see Figure 5 and Appendix 6 for details). When the *α* parameter is 1, the topology realizes Weismannian inheritance, and the diversity of X sequences in the population is at its possible minimum (Figure 5). As *α* decreases towards 0, inheritance becomes more and more Lamarckian, *y*_*t*_ defining *x*_*t*+1_ more and more, and the diversity of X sequences increases to the maximum (or to the population size, if that is the smaller limit; Figure 5).

**Figure 5.**
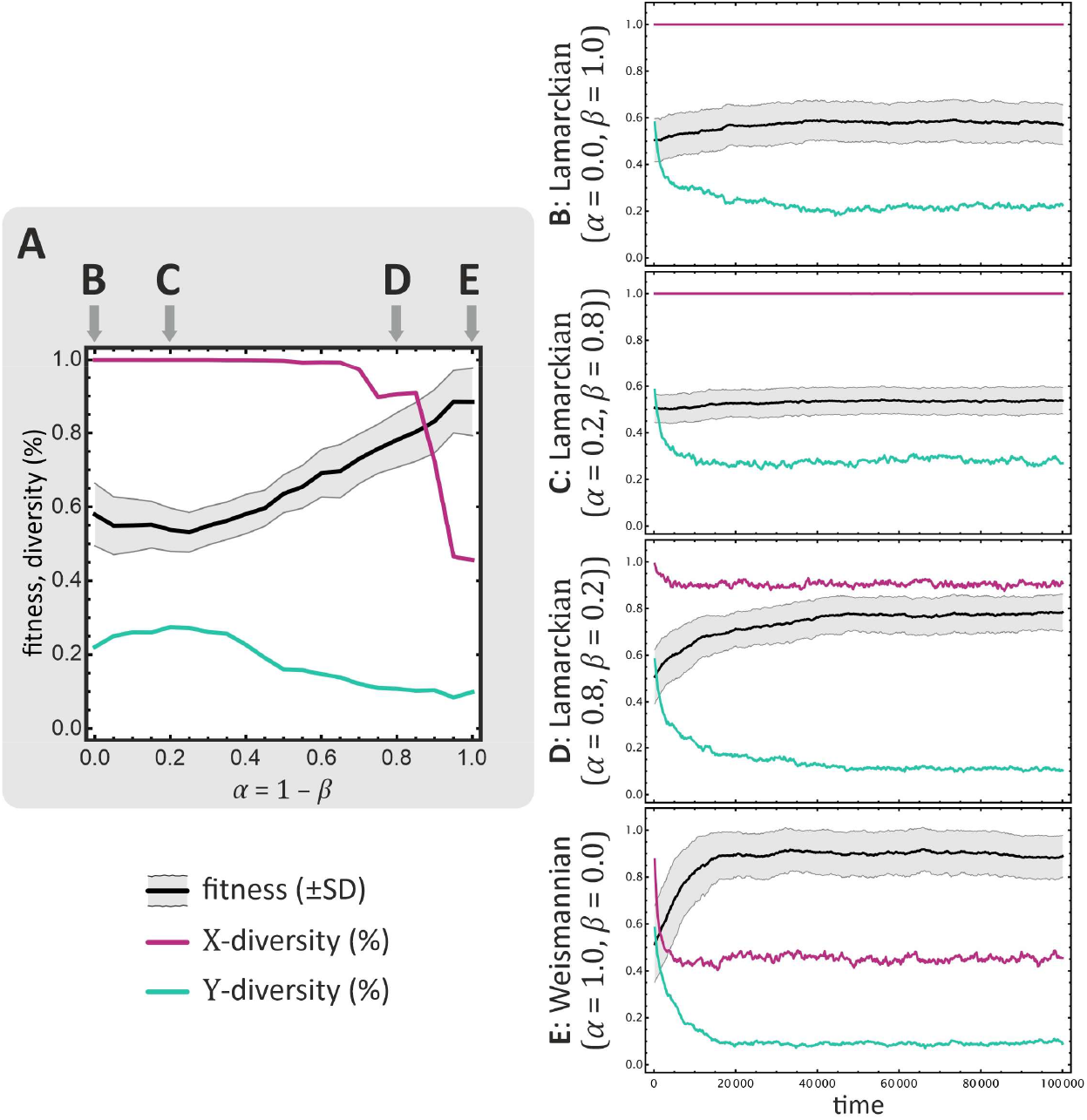
Comparison of replication-selectional dynamics of Weismannian and Lamarckian populations. At each timestep, a single sequence is selected for replication from the population with probability proportional to its fitness. Fitness is the linear combination of w(x_t_) and w(y_t_), according to the mixing parameter β; the next generation x_t+1_ is created from {x_t_, y_t_} according to the mixing parameter α (cf. Figure 4G). During replication, mutations can happen with per-digit probability μ in any transformation. Thick black curve shows mean fitness with ± SD values, purple and teal curves show X and Y sequence diversities, respectively (i.e. the number of different types of sequences of state X and Y, as percentage of the population size). **A:** Crossing the continuum of Lamarckian-Weismannian replication at α = 1 − β (diagonal of Figure 6). Each datapoint shows the average of the equilibrium of a simulation with a specific {α, β} pair. **B-E**: Individual simulations of plot **A. B**: Simultaneous Lamarckian inheritance (Figure 4G). **C, D**: Successive Lamarckian inheritance (Figure 4F). **E**: Weismannian population (Figure 4C). Parameters are: N_X_ = 30, N_Y_ = 10, c_X_ = 3, c_Y_ = 1, a_X_ = a_Y_ = 2, θ_XY_ = θ_uniformly ordered_, 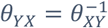, μ_XX_ = μ_XY_ = μ_YX_ = 0.01, population size of 1000, and 100 000 replication steps. For more information, see Appendix 6.

Analysing the model of 2-state informational topologies, it becomes clear that ultimately, there are only three formally distinct informational topologies for 1 or 2 distinct states that, in principle, can maintain information for an indefinite time (Figure 4). We call these *direct, Weismannian* and *Lamarckian* inheritance, corresponding to the simplified schemes of Figure 3. Other possible topologies are either isomorphic to one of the three or are incapable of maintaining information for an indefinite time. According to our formalism, Lamarckian replication corresponds to probabilistic replication of the first type (Figure 4D), where the intermediate external state *y*_*t*_ is directly generated from the internal state *x*_*t*_ and is also directly responsible for the next generation of internal state *x*_*t*+1_. Contrarily, in Weismannian replication, the external *y* state (soma) never contributes to the next generation of *x* (germline) directly.

### 3. Replication topologies form a continuum

Based on the above observation that the functional division of labour need not be absolute with exclusively dedicated states, we conclude that the three topologies form a continuum (Figure 6). Direct (Figure 4A) and Weismannian replicators (Figure 4C) constitute the extrema of this continuum. In Weismannian inheritance, there exist dedicated states for information inheritance and interfacing selection; in direct replication, there is a single state responsible for both roles (additional states may exist, with no role in selection/evolution). Contrarily, in Lamarckian inheritance (Figure 4D-G) Figure 4D-F, corresponding to antidiagonal corners inFigure 6Figure 4Figure 6. Lamarckian replication differs from direct replication in that it has two distinct states, similarly to Weismannian inheritance. However, unlike in Weismannian inheritance, heritable information is not protected within a germline but is exposed as an external intermediate before being transformed (fully or partially) to internal state again. That is, functions of conserving information and interfacing selection are not entirely separated (similarly to direct inheritance). This justifies the existence of the third type, besides direct and Weismannian replication.

**Figure 6.**
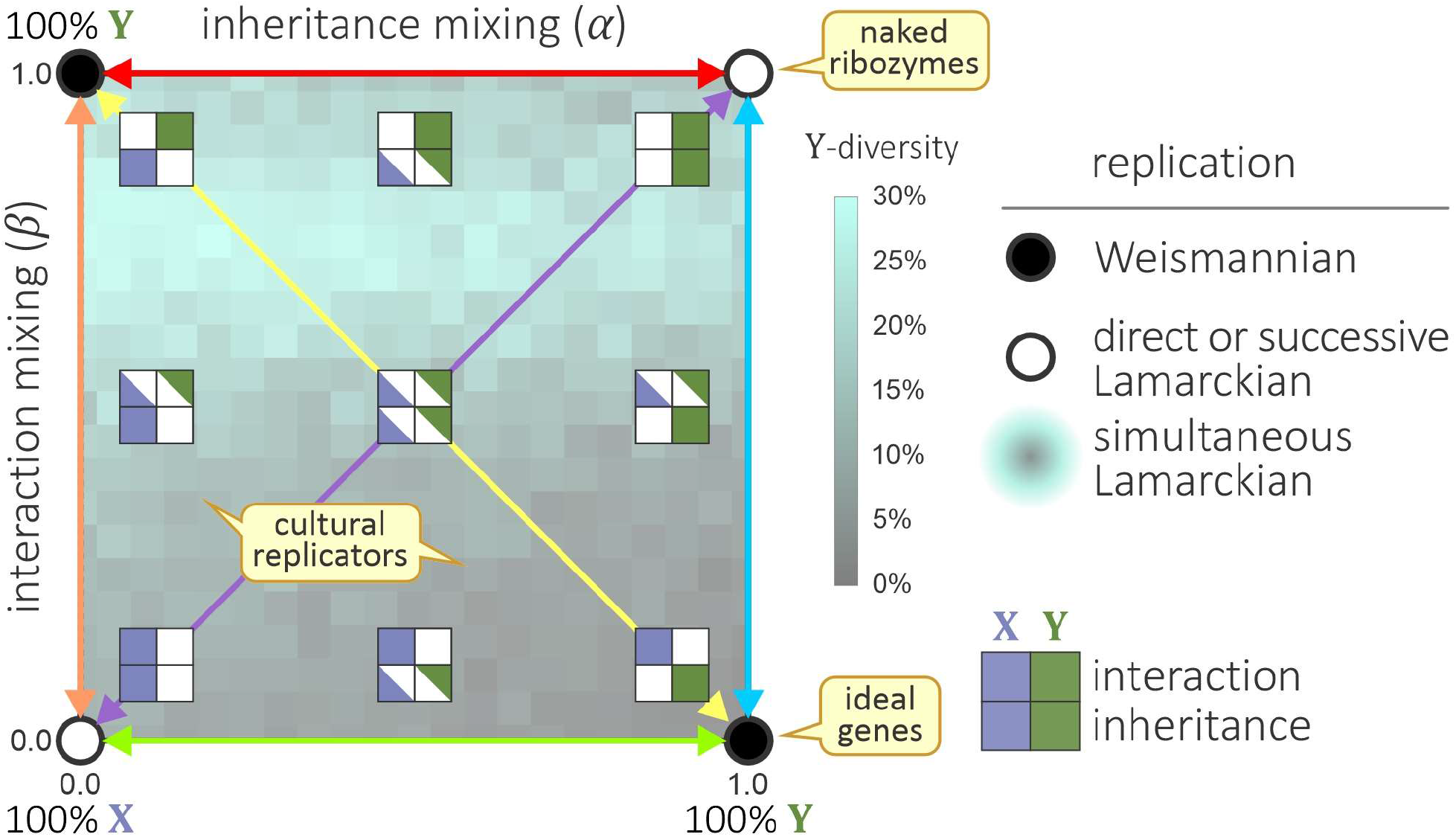
The functional continuum of 2-state informational inheritance systems. Inheritance systems involve up to 2 states, x ∈ X (purple) and y ∈ Y (green), responsible for two biofunctions, maintaining information for an indefinite time (inheritance) and being subject to direct selection (interaction with environment). Empty circles represent direct replication systems where only one state is responsible for both roles (Figure 4A, D-F), filled circles represent ideal Weismannian inheritance systems (Figure 4C). The farther from the corners the more pronounced the distributed nature of biofunctions becomes. In Weismannian inheritance (Figure 4C), Y is responsible solely for information storage while X is responsible for interaction, i.e., Y is a perfect germline and X is a perfect soma. Any system not at the corners is by definition Lamarckian (Figure 4G). The background represents the actual Y diversity according to simulations in Appendix 6. The colored arrows indicate simulations represented in Figure SI 11, red arrow corresponds to Figure 5. Opposite corners are formally isomorphic up to a replacement of X ↔ Y, therefore the continuum can be mirrored at both diagonals.

Presumably, inheritance systems can gradually transform to each other via evolution within this continuum, that is, a Lamarckian system may evolve toward Weismannian inheritance, toward being more effective in conserving information on a longer timescale. It is a question of actual material realization of the replicator and interactor whether this evolutionary transformation is possible or not.

### 4. Lamarckian inheritance maintains the information of the external state

The distinct topology of Lamarckian inheritance (Figure 3C) has some interesting properties. It is true, that the replicator *x*_*t*_ is the template of the interactor *y*_*t*_, but the interactor *y*_*t*_ is also the template of the next generation of replicator*x*_*t*+1_, unlike in Weismannian replication. Since in culture, the direct transformation X → X does not exist (no direct template copying from brain to brain), the interactor *y* becomes an equally valid replicator as *x*.

Due to the lack of direct template copying, there is a potential source of information loss inherent to the replication mechanism if any of the transformations is a many-to-one mapping (see Figure SI 1). This means that during this transformation, some information is always lost. It is possible, that all transformations involved in a Lamarckian topology are bijective (all are transcriptions, Figure SI 1), in which case inheritance can maintain the information encoded in *x*_*t*_ (and *y*_*t*_) indefinitely. However, whenever there is non-invertible transformation *θ*_T_: X → Y (as in Weismannian inheritance from germline to soma), compression cannot be unambiguously inverted to reconstruct the original state, and Lamarckian topology loses information of *x*. That is, the lack of template copying likely entails that the switching between internal and external states loses information during transmission **even if there is no exogenous error** during inheritance (see Figure SI 8).

However, this inherent vulnerability to informational loss does not mean that the Lamarckian topology cannot maintain information. If we assume that the switch from internal to external state (*θ*_XY_: X → Y) reduces the representational space (i.e. *θ*_XY_ = *θ*_T_), then the process loses information in the transformation because different *x*-s are mapped to the same *y*. Nevertheless, Lamarckian inheritance can maintain information stably, if the reverse of *θ*_XY_ is a mapping *θ*_YX_: Y → X that can stably yield *x*_*t*+1_⬚ **that is equivalent under selection** to*x*_*t*_, otherwise the whole process is not replication anymore. That is, if 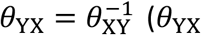 is the probabilistic inverse of *θ*_XY_, see Appendix 1), inheritance can maintain information, but only as much as is encoded in the state with the smaller cardinality (in this example, Y, the image of *θ*_XY_). Thus, **Lamarckian inheritance can only maintain as much information as is represented by the external state**. The argument is the same for the isomorphic case where roles of *x* and *y* are switched.

Accordingly, we distinguish three different sources of information loss, where original information is replaced with something else:

1. **Exogenous error**: Random environmental factors perturbing transmission channels or states. Analogous to e.g., point mutation of DNA caused by radiation, or noise pervading a communication channel in culture.
2. **Many-to-one transformation**: The nature of the transformation *θ*: X → Y is such that it maps to an image of reduced size or dimensionality. A successive transformation *θ*^-l^: Y → X that is the (probabilistic) inverse of the first one may return a selectively equivalent state *x*_*t*_*∼x*_*t*+1_, ∀*t*, but the information content that can be maintained is reduced (see Figure SI 2 and Figure SI 3). Presumably, development works like this, where the genetic information is translated to protein-space. In culture, production (externalization of mental representation) is bound to reduce information content.
3. **Asymmetric transmission**: Information is lost due to successive transformations not being (probabilistic) inverses of each other ((*θ*_*t*_)^*-l*^ ≠ *θ*_*t*+1_ in Figure 4). In this case, not even selectively equivalent states can be maintained. Replication becomes severely limited, or attractor-based (cf. ^116^).

That is, if *θ*_YX_ is not the probabilistic inverse of *θ*_XY_, then *θ*_YX_ cannot counter translation *θ*_XY_, and loss of information is inevitable, leading to the accumulation of errors over time. The same fate does not apply to Weismannian topology, as the state *y* (soma) can be arbitrarily reduced. Since *y* never contributes to the next generation of *x* (genotype) during replication, this reduction need not be countered to maintain the information content of *x*. An information losing transmission, strictly speaking, is not informational replication, as it cannot stably maintain hereditary information indefinitely.

### 5. Lamarckian inheritance can approximate Weismannian inheritance

Genetic replication relies on template copying, but in general, a replication process can be composed of a sequence of arbitrary transformations, as was explained above (Figure 4). Calculations of the replication fidelity of the Lamarckian inheritance method show that Lamarckian replication can be exact, even in case of probabilistic transformations and a pair of successive transformations that are probabilistic inverses of each other can approximate the copying fidelity of Weismannian replication (Appendix 3 and 4). A probabilistic transformation (e.g. *θ*_XY_) in a replication process therefore does not necessarily cause replication to lose information: if it is accompanied by the correct inverse transformation 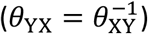, the probabilistic effect can be cancelled. Weismannian and Lamarckian replicators thus can maintain identical amount of information concerning Y-sequences, though Lamarckian inheritance maintains a larger X-diversity (see Figure 5 and Appendix 5). The point is, that if an inheritance method and topology can stably maintain information throughout successive transformations, it implements replication, regardless of whether it relies on bitwise template copying or other mechanisms. It becomes a quantitative matter how well cultural internalization and reconstruction (Lamarckian Y → X) approximates the inverse of externalisation (Lamarckian X → Y). The mechanisms involved in cultural (and language) inference and reconstruction to minimize such errors are manifold (see Discussion). We point out that these steps do not counter the argument that cultural evolution is Darwinian on the population level ^27–30,32,33^. In fact, they effectively yield a Darwinian process despite Lamarckian inheritance underlying it (cf. Figure 7).

**Figure 7.**
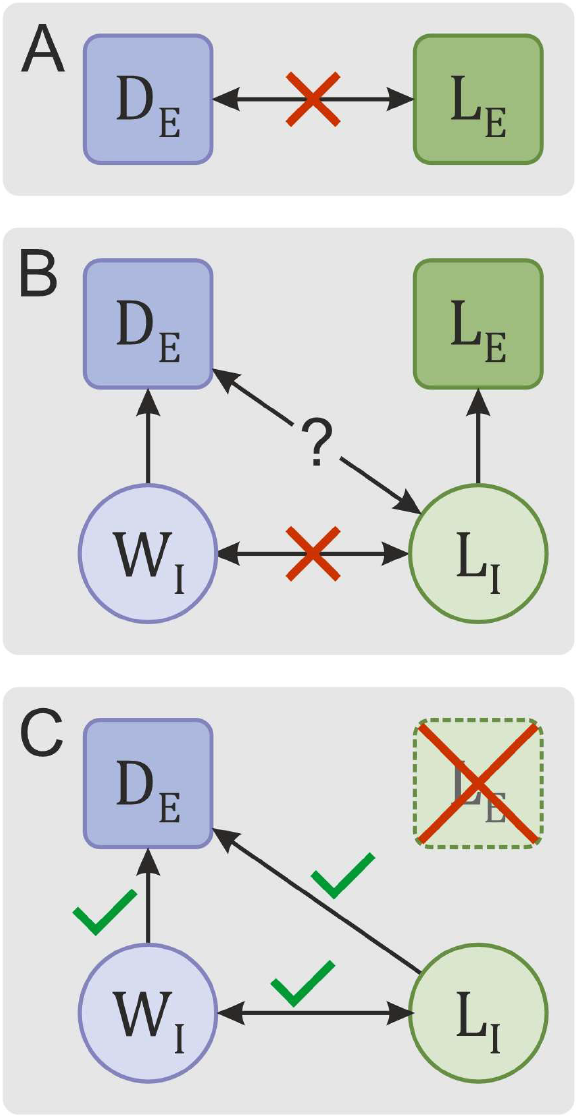
The relation of Darwinian and Lamarckian inheritance, in historical context. **A**: Darwinian evolution was traditionally pitted against Lamarckian evolution. Lamarck claimed that use and disuse (guided variation) change biological functions and individuals, and it is the inheritance of these changes that drives Lamarckian evolution (L_E_). This contradicts Darwin’s claim that variation is blind and it is natural selection acting on variation that drives Darwinian evolution (D_E_). **B**: According to a common interpretation, Lamarckian inheritance (L_I_) does not involve guided variation but still accounts for the inheritance of lifelong changes. Therefore, it is incompatible with August Weismann’s insight that it is the germline that exclusively inherits information between successive generations (Weismannian inheritance, W_I_). **C**: In our interpretation, Weismannian and Lamarckian inheritance are part of the same continuum, and their difference is quantitative (how much the separate states divide labour). Accordingly, in theory, both can yield Darwinian evolution.

## Discussion

We have reconciled the different inheritance methods, Lamarckian and Weismannian, on a formal basis. We have demonstrated that Lamarckian and Weismannian inheritance are part of the same continuum of informational topology. There are actually three distinct topologies that are replicative in nature: direct, Weismannian and Lamarckian inheritance (Figure 3). Direct replication corresponds to e.g., naked ribozymes replicating, without a vehicle or a distinct protein shell to provide a distinct phenotype. Weismannian inheritance corresponds to how biological organisms inherit information through the germline, and face selection through the soma. Lamarckian inheritance corresponds to cultural inheritance. The Weismannian and Lamarckian topologies can only be defined if two distinct and at least partially separate states exist in the life cycle of inherited information: the germline and soma ^79^ or internal and external cultural items ^68^, corresponding to the replicator and interactor *in abstracto* ^119^. This distinction is only possible if phenotypic changes are “stored” elsewhere than genotypic changes.

Our model naturally provides a unification of these inheritance topologies as a continuum, under the assumption that the two states may gain or lose dominance in conserving information and/or interfacing the selective environment (Figure 6). In cultural inheritance, the external product (utterance, artefact) mediates information transmission between brains and is directly transformed to the new internal representation (mental state). As a result, both states may inherit acquired changes. Lamarckian replication thus differs from direct replication in that it has two distinct states, one being a dedicated interactor, similarly to Weismannian inheritance. However, unlike Weismannian inheritance, the function of conserving information may not have a dedicated state. In this continuum, direct and Weismannian replicators signify the far ends while Lamarckian replicator is anything in-between the two extrema. We therefore propose the following definition of Lamarckian replication:

***Lamarckian replication*** *is an informational replication process of an inheritance system with at least two separate states of heritable information, but without fully separate roles of storing heritable information and interfacing selection*.

A change in the informational topology itself does not mean that Lamarckian inheritance is inferior to Weismannian regarding replication. Both topologies could, in principle, rely on transformations that do not lose information at all. However, the snag is, that neither the transformation of genotype to soma in Weismannian inheritance, nor the production of cultural items from mental states can be trivially inverted. That is, information is inevitably reduced when switching from one state to the other (e.g. developing the soma or producing an utterance). This does not pose an issue for Weismannian inheritance, where the soma is not expected to contribute considerable heritable information to the next generation. However, in Lamarckian inheritance, the “template” for the next generation is the externalized item (e.g. utterance), which may entail an ultimate loss of information.

We have demonstrated that, coupling appropriate inverse transformations can ensure that there is no information loss in an ideal situation (without mutation). In theory, all states of a Lamarckian topology are equivalent under selection. This means that items of **the internal and external culture act as templates of each other, representing the same information in different media, and collectively functioning as a unit of replication**. Practically speaking, all states replicate, and they can stably maintain the same amount of information. However, this amount is defined by the state with the smallest representation space (likely the external one).

Thus, replication need not rely on direct template copying (as in DNA replication) to inherit and stably maintain information, at least in principle. As we have demonstrated, the copying fidelity of Lamarckian replication can approximate that of Weismannian inheritance and can support Darwinian evolution, even under a nonzero mutation rate. It also means that the same replication dynamics and fidelity can emerge from different inheritance topologies (see e.g. ^28,45^). Lamarckian inheritance can maintain the same amount of information as Weismannian inheritance, though Lamarckian inheritance maintains larger diversity of internal states (see Appendix 6). A large neutral diversity may be beneficial, for example in case of rapidly changing environments ^120^, but also means a larger mutational kernel and load.

Culture (and cultural agents, genetically) likely evolved to compensate for the excess neutral diversity of internal states and the excess/contradicting variability introduced by multiple parents and channels. Culture entertains many mechanisms that increase the replication fidelity of inheritance. These mechanisms ensure that despite the Lamarckian topology and potentially conflicting sources, inheritance approximates Weismannian fidelity as closely as possible (allowing us to apply our template-formalism to culture). Without an exhaustive enumeration, we mention the followings: repetition of examples ^34,121,122^ and metacommunication can reduce ambiguity ^123,124^; composition of partial information from multiple sources ^46^ and a statistically regular linguistic structure can help reconstruct incomplete information ^113^ e.g. by compositional and regular languages ^34,125^; the transmission of generative rules (e.g. recipes) instead of mimicking (copying examples directly) ^38,59,76,126^; persistent artefacts, codification, stable references and digitalization can preserve information over generations ensuring high copying accuracy (tools, steles, codices, books, blueprints, repositories, the internet, etc.) ^29,127–130^; evolved cognitive functions (deduction, the ability to model other individuals’ point of view) can be used to find missing information (e.g. context) or to correct errors.

Cavalli-Sforza and Feldman have investigated the population genetics of cultural inheritance by a set of successively refined models ^44,89–94^. They considered a population of social organisms with a diallelic diploid ^90^ or haploid ^92^ representation of heritable information, two genes, one passed on genetically, one culturally to the offspring. Given a transmission matrix that defines the transmission rates of geno-phenotype combinations to each other (plus mutation, selection and population structure), they applied equilibrium theory to investigate whether a stable polymorphic population can exist where all geno-phenotype combinations survive ^92^.

They assumed continuous traits and explored various scenarios where genetic and cultural traits affect (or not) each other’s inheritance. They have separately considered *i*) mutation, *ii*) transmission source (population structure and cultural parents), and *iii*) selection (differential survival of the various geno-phenotype combinations). They have explored a range of models between complete genetic control and extreme environmental (cultural) determination, investigating the change in frequencies over time, concluding that “between the two extremes is a continuum of intermediate models” ^93^, resonating with our assessment of the Lamarckian-Weismannian continuum.

Their models measure the correlations between biological relatives based on the amount of phenotypic plasticity (environmental/cultural influence) but over the distribution of phenotypes in the population ^93^ and not over the amount of information that can be maintained by the transmission channel *per se*. These two are complementary. Later, they have extended the model to include a temporally changing transmission matrix, considering some basic matrix types ^89^ and to study discontinuous phenotypes ^90^, while ultimately arriving to a haploid two-gene model ^92^. Regarding their transmission matrix parameters, they write: “The transmission coefficients *β*_*i*_ and *γ*_*i*_ are probabilities that describe complicated processes by simple numbers. They are intended to represent relative values of the rate at which one or other state can be achieved via observation, imitation, and education” in cultural inheritance ^92^. Our model can be understood as the extension of the model of Cavalli-Sforza & Feldman by providing a method to calculate key parameters: transmission rates *γ* and *β* (see ^92^).

We believe that the two aspects, information theory of underlying mechanisms and population genetics can be combined to give a fuller explanation of cultural evolution. This is left for future work though.

## Conclusions

According to universal Darwinism, evolution is a domain-independent process that emerges whenever the criteria of evolutionary units are met (multiplication, heredity and variability ^131^). These units may be responsible for non-genetic adaptive systems, like culture ^27^. Whenever an inheritance system can stably reconstruct and maintain information throughout successive generations, it implements replication, regardless of whether it relies on template copying or other means. Socially transmitted traits (e.g. linguistic, religious or political) show remarkable stability while maintaining variation ^85^, attesting to the evolutionary requirements of Darwinism.

However, due to the lack of a digital structure, template copying and a proper division of labour between informational states, cultural replication is often termed Lamarckian ^13,26,38,74^ or not even considered replication at all, despite clearly showing Darwinian properties. This duality of cultural inheritance poses an apparent paradox. However, the lack of discrete self-replicating digital units at the lowest level (memes akin to genes) does not disqualify Darwinian evolutionary models from the cultural domain ^28,84,109,118^.

Here we show that there is no paradox between the micro-level observations suggesting Lamarckian inheritance and the macro level properties suggesting Darwinian evolution. We show that **an inheritance method that can inherit acquired changes (i.e. is Lamarckian) can nevertheless maintain Darwinian evolution**. Moreover, we show that under certain conditions the fidelity of Lamarckian inheritance (strictly speaking, the amount of information that can be stably maintained without selection) can approximate that of a Weismannian system.

The fact that both Weismannian and Lamarckian inheritance can maintain Darwinian evolution accounts for the apparent duality of culture. We believe that there is no contradiction between the claims that culture is Lamarckian on the micro-level, yet Darwinian on the macro-level. According to our understanding, **cultural inheritance is inherently Lamarckian locally, while cultural evolution is Darwinian on the population level**. Lamarckian inheritance, due to its lack of template copying, might inherently lose information at individual transmission events (locally), but iterative and inferential mechanisms are often utilized to average over multiple sources, lineages and iterations to approximate template copying (at the population level). As a result, cultural inheritance is mostly Lamarckian in mechanism but is Darwinian in its effect (Figure 7).

Lamarckian inheritance is different from Weismannian in three aspects: 1) it inherits changes acquired by the soma (interactor, external culture); 2); it likely involves transformations that reduce representations; and 3) it may involve non-invertible transformations, cumulatively decreasing replication fidelity. As a result, Lamarckian inheritance can only maintain as much information as the soma, effectively making the soma the real genotype of Lamarckian replicators. Formally, Lamarckian inheritance is reduced to Weismannian inheritance with low fidelity, where the genotype is actually not stored exclusively by the internal state but by the internal and external states jointly. In summary, Lamarckian inheritance is consistent with Darwinian evolution, and it becomes a quantitative matter to what extent soma-acquired changes are inherited or discarded.

The problem of the critiques of potentially Darwinian culture (and memetics ^30,60,132^) is that no replicating particles seem to exist for which Darwinian dynamics can be applied, and therefore “there is no guarantee that the mental representation in the second brain is the same as it is in the first” ^28^. We have explicitly considered these concerns in our paper. We believe that no particles, nor strict identity is necessary to maintain information persistently via Darwinian replication. As our model proves, it is *information* that must be replicated to maintain culture, and physical objects are as much the vehicles of this information as social organisms and the environment. Darwinian dynamics is not about replicating matter but about replicating information (even if it is attractor-based ^28,45^ and not storage-based like DNA ^4,116^), as was clearly demonstrated by e.g. Eigen’s formal model of informational replicators ^117^. Moreover, as we have proven, successive cultural replicators need not be identical while they code for the same thing (i.e. are phenotypically equivalent under selection).

To explain the stability of cultural inheritance, one need not expand evolution to multiple dimensions ^13,133^ or introduce purported new mechanisms (see ^134^ and its critique ^135^). We have demonstrated that informational replication in biology and culture, though quantitatively differing, can be approximated by and unified within the same formalism of template-based information transmission, complementing the foundational models of cultural evolution ^90,92,93^. Despite the fact that culture implements Lamarckian inheritance, it can nevertheless yield Darwinian evolution. Only the continued selection of Darwinian replicators can maintain adaptive evolution against entropy.

## Methods

We formally describe an inheritance system as an informational topology involving up to two distinct information-bearing (i.e., digital), persistent states, *x* ∈ X and *y* ∈ Y. Heritable information is stored by these states and inheritance is modelled as transmission of information between successive states, represented by a chain of **transformations** *t*_*l*_, *t*_2_, *…* mapping between states X and Y. In this paper, we only consider **compositional transformations** *T*_*i*_, i.e. transformations for which an **elementary transformation** *Θ*: *S*_X_ → *S*_Y_ (or *Θ*: *S*_Y_ → *S*_X_) exists such that:

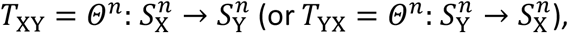

where (*n*∈ℕ) ≥ 1 is the number of modules (e.g., codons) and *S*_X_ ⊂ X and *S*_Y_ ⊂ Y are the module domains (e.g., codon sets) of sets X and Y, respectively. That is, an elementary transformation maps codon-spaces (e.g., from RNA codons to amino acids), a compositional transformation maps polymer sequence-spaces.

TThe elementary *θ* may be many-to-one (forward translation *θ*_*t*_, where image is smaller than domain), one-to-many (backward translation *θ*_R_, where image is larger than domain) or maintain domain cardinality (transcription *θ*_S_, or copying *θ*_C_, if domain and image are identical; see Figure SI 1). We assume, that the *θ* are deterministic, mutation not being inherent to them (see Appendix 2 for further model details). We do consider mutation in the full analysis (see Appendices 3-6), where mutational operators *m*_*ij*_ are applied to the result of each transformation, e.g., *y*_*t*_ = *m*_XY_(*θ*_XY_(*x*_*t*_)), meaning that the result of translation *θ*_XY_(*x*_*t*_) is mutated with per-digit mutation rate of *μ*_XY_ specific to the transformation *θ*_XY_ to yield actual *y*_*t*_.

During **complex information transmission** Θ, elementary transformations are chained (in succession, Figures 2, 3; or in parallel, Figure 4G):

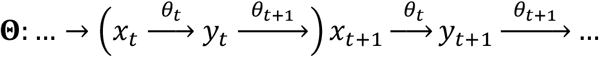

where *x*_*t*_ and *y*_*t*_ represent states at time step *t, θ*_*t*_ = *θ*_XY_ and *θ*_*t*+1_ = *θ*_YX_. Note, that parentheses denote the repetitive subset of steps we actually model. In culture, we assume, that *x* is the internal and *y* is the external state, noting that these terms are only important for illustrative purposes and bear no formal relevance. The important point is that we assume that changing from the internal to the external state by *θ*_XY_ may involve loss of information that *θ*_YX_ might not be able to reconstruct. When *y* is not involved in generating *x*, the route of information inheritance simplifies to *x*_*t*_ → *x*_*t*+1_ → x_*t*+2_ → …, corresponding to direct or Weismannian replication (Figure 3A orB).

For Θ to maintain evolutionary information potentially indefinitely, the minimal condition is that **successive states must be equivalent under selection**. *x*_*t*_∼*y*_*t*_∼*x*_*t*+1_∼*y*_*t*+1_∼ …, ∀*t*. Sequences *x*_*i*_, *x*_*j*_ ∈ X or *y*_*i*_, *y*_*j*_ ∈ Y are **phenotypically equivalent** (∼) if they share the same fitness under selection:

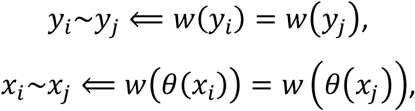

where *w* is the fitness function defined over Y, representing the distinct phenotypes of elements of X under selection. (for details, see ^4^). Relying on equivalency instead of structural identity allows other transformations than template copying to be involved in the replication process. However, with elementary transformations that do not map information one-to-one, information can be potentially lost, specifically if *x*_*t*_ ≁ *x*_*t*+1_ or *y*_*t*_ ≁ *y*_*t*+1_. If only one of the conditions is true, the system can only maintain as much information as the upholding condition allows (the equivalence on *x*_*t*_ *or* on *y*_*t*_). For more details, see Figure 4 and Appendix 2.

However, if the successive elementary transmissions are (probabilistic) inverses of each other, a certain amount of information can stably be maintained even if elementary transformations are not 1-to-1. The **probabilistic inverse** of *θ*_*t*_ is defined as:

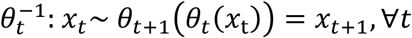

that is, *θ*_*t*+1_ is the probabilistic inverse of *θ*_*t*_ if *x*_*t*+1_ is phenotypically equivalent to *x*_*t*_ after 2 successive transformations by *θ*_*t*_ and *θ*_*t*+1_. This is trivially true if *θ*_*t*_ and *θ*_*t*+1_ are template copying, but not necessarily if any of them is a forward or backward translation (see Figure SI 1). This is likely the case in culture, where information is reductively translated from the internal level to the external level, and the external item serves as a template for the next generation. Consequently, during an information transmission process, information can be lost either due to random mutation posed by external factors or inherently, due to the informational topology of inheritance, when a successive pair of elementary transmissions are not the (probabilistic) inverses of each other. Accordingly, we emphasize, that for replication to happen, it is not necessary that *x*_*i*_ = *x*_*j*_ or *y*_*i*_ = *y*_*j*_ for all *i* ≠ *j*.

We have derived the amount of information that can be maintained in case of Weismannian and Lamarckian information inheritance topologies, under similar external mutation rates (Appendices 3, 4, respectively). We have statistically measured these values (Appendix 5) and performed population-based simulations (see Figure 5 and Appendix 6), to confirm our results.

## Supplementary Material

## 1. The shared vocabulary of biological and cultural replication

**Table SI 1.**
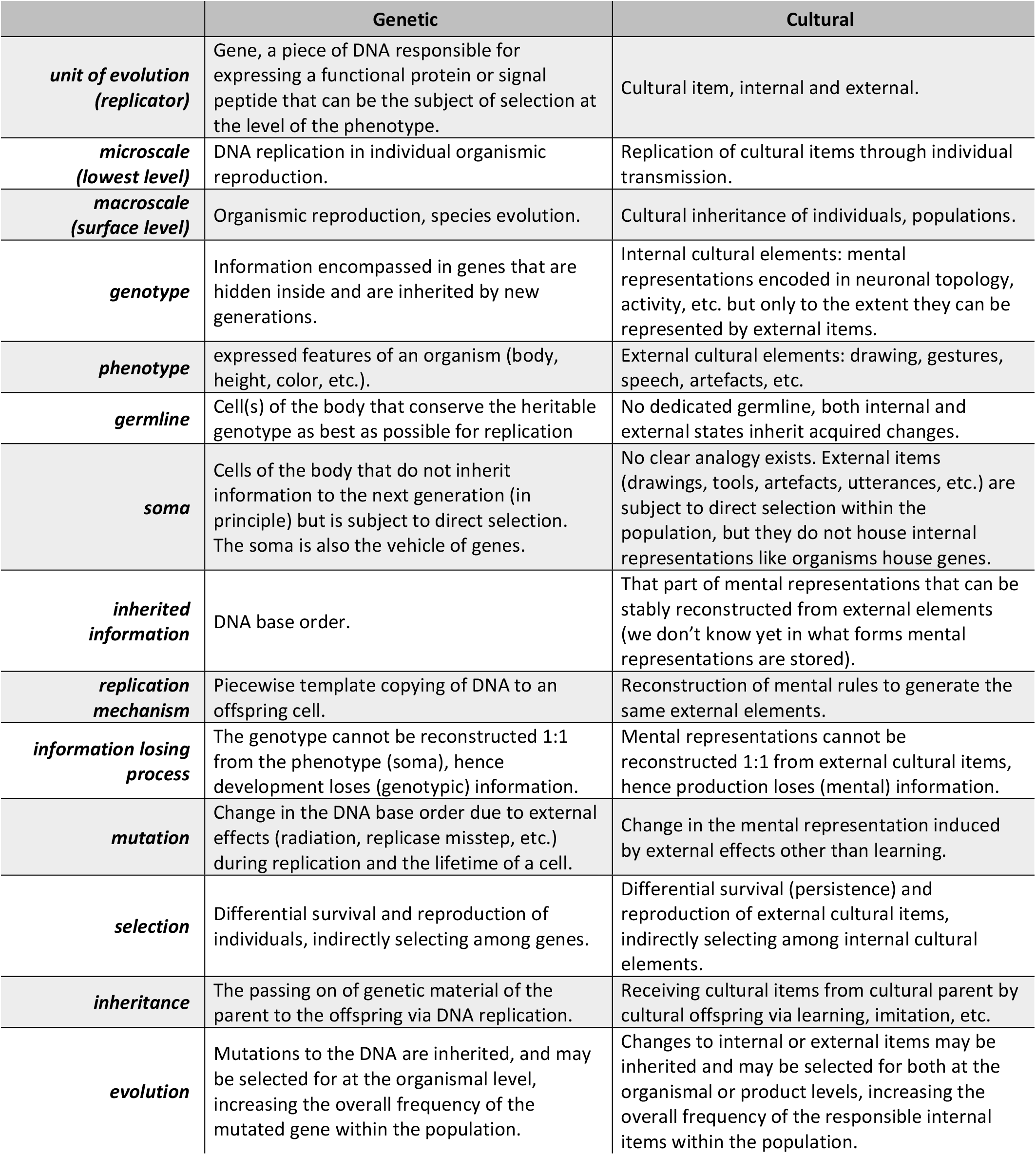
The shared vocabulary of biological (genetic) and cultural information replication, as a reflection to the lack of common language emphasized by ^64^.

|  | Genetic | Cultural |
| --- | --- | --- |
| **unit of evolution (replicator)** | Gene, a piece of DNA responsible for expressing a functional protein or signal peptide that can be the subject of selection at the level of the phenotype. | Cultural item, internal and external. |
| **microscale (lowest level)** | DNA replication in individual organismic reproduction. | Replication of cultural items through individual transmission. |
| **macroscale (surface level)** | Organismic reproduction, species evolution. | Cultural inheritance of individuals, populations. |
| **genotype** | Information encompassed in genes that are hidden inside and are inherited by new generations. | Internal cultural elements: mental representations encoded in neuronal topology, activity, etc. but only to the extent they can be represented by external items. |
| **phenotype** | expressed features of an organism (body, height, color, etc.). | External cultural elements: drawing, gestures, speech, artefacts, etc. |
| **germline** | Cell(s) of the body that conserve the heritable genotype as best as possible for replication | No dedicated germline, both internal and external states inherit acquired changes. |
| **soma** | Cells of the body that do not inherit information to the next generation (in principle) but is subject to direct selection. The soma is also the vehicle of genes. | No clear analogy exists. External items (drawings, tools, artefacts, utterances, etc.) are subject to direct selection within the population, but they do not house internal representations like organisms house genes. |
| **inherited information** | DNA base order. | That part of mental representations that can be stably reconstructed from external elements (we don't know yet in what forms mental representations are stored). |
| **replication mechanism** | Piecewise template copying of DNA to an offspring cell. | Reconstruction of mental rules to generate the same external elements. |
| **information losing process** | The genotype cannot be reconstructed 1:1 from the phenotype (soma), hence development loses (genotypic) information. | Mental representations cannot be reconstructed 1:1 from external cultural items, hence production loses (mental) information. |
| **mutation** | Change in the DNA base order due to external effects (radiation, replicase misstep, etc.) during replication and the lifetime of a cell. | Change in the mental representation induced by external effects other than learning. |
| **selection** | Differential survival and reproduction of individuals, indirectly selecting among genes. | Differential survival (persistence) and reproduction of external cultural items, indirectly selecting among internal cultural elements. |
| **inheritance** | The passing on of genetic material of the parent to the offspring via DNA replication. | Receiving cultural items from cultural parent by cultural offspring via learning, imitation, etc. |
| **evolution** | Mutations to the DNA are inherited, and may be selected for at the organismal level, increasing the overall frequency of the mutated gene within the population. | Changes to internal or external items may be inherited and may be selected for both at the organismal or product levels, increasing the overall frequency of the responsible internal items within the population. |

## 2. Model definition

### Transformations

We define **transformations** *t*_XY_: X → Y as a mapping from set X to set Y, and *t*_YX_: Y → X as a mapping from Y to X, for all *x* ∈ X and *y* ∈ Y:

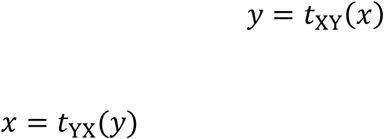

A transformation describes how e.g., RNA sequences (X) map to poly-amino-acids (Y). Certain transformations can be represented by simpler transformations, that consistently map shorter parts of sequences to their correspondents, e.g. codons to amino acids. Therefore, we define a **compositional transformation** *T*_XY_: X → Y (and *T*_YX_: Y → X) as a mapping for which an **elementary transformation** *Θ*_XY_: *S*_X_ → *S*_Y_ (and *θ*_YX_: *S*_Y_ → *S*_X_) exists such that:

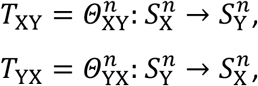

where (*n*∈ℕ) ≥ 1 is the number of modules (e.g., codons) and *S*_X_ ⊂ X and *S*_Y_ ⊂ Y are the module domains (e.g., codon sets) of sets X and Y, respectively. The sizes of the codon domains are defined as:

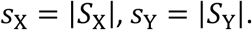

Two sequences *x*_i_, *x*_*j*_ ∈ X or *y*_i_, *y*_*j*_ ∈ Y are defined to be **phenotypically equivalent** (∼) if they share the same fitness under selection:

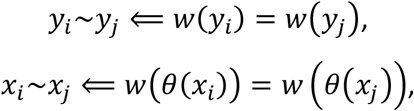

where *w* is the fitness function defined over Y, representing the distinct phenotypes of elements of X under selection. In other words, two sequences *x*_*i*_ and *x*_*j*_ are phenotypically equivalent if selection handles their phenotypes identically in a short timeframe (i.e. in a single selection event, to omit e.g. the different mutation kernels available for different genotypes). For the sake of generality, we do not specify selection and fitness; the point of introducing phenotypic equivalency is to ensure that sequences can be different in their genotypes (e.g. *x*_l_ and *x*_2_) while still have the same phenotype (*θ*(*x*) ∈ Y). For consistency, we denote the fitness function as *v*, when applied directly over X, noting that it yields the same partitioning of Y under selection as *w* does over *θ*(*x*), ∀*x* ∈ X, under the same selection:

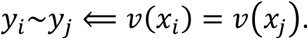

We now can meaningfully define the **probabilistic inverse** *θ*^-l^ of a transformation *θ* as a transformation that, when applied to *θ*(*i*), yields sequence *j* which is phenotypically equivalent to *i*:

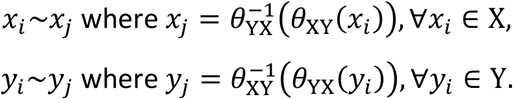

This is trivially true if *θ* and *θ*^−l^ are one-to-one mappings, but not necessarily if any of them is not (assuming no mutation, see Figure SI 1). For sake of simplicity, we use familiar terms of polymer processing:

- *θ*_*t*_(**forward translation**) is a many-to-one function (i.e., surjective, but non-injective function). Genetic translation maps codons to amino acids in a way that for every amino acid, there is at least one codon that maps to it. Such a function cannot be inverted without using probabilistic rules to map back elements of the image to the domain (see next).
- *θ*_R_ (**backward translation**) is the inverse of a surjective, but non-injective mapping, hence is a multivalued one, mapping one-to-many in a probabilistic nature. We consider only cases where possible multiple values are selected with equal probabilities. Such a transformation is, by definition, ambiguous, as multiple distinct elements of the image represent the same element of the domain. A hypothetical example is the probabilistic reverse translation of amino acids to polynucleotide codons.
- *θ*_S_ (**transcription**) is a bijective mapping between potentially non-identical domain and image, hence is a one-to-one function. Applying the function twice should not change the argument. For example, genetic transcription maps DNA nucleotides to RNA nucleotides in a one-to-one method.
- *θ*_C_ (**copying**) is a special case of transcription where domain and image of the mapping are identical, thus *θ*_C_ is the identity function, for which it is always true that 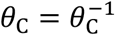. For example, template copying of DNA replication ensures that the same nucleotide order appears, though it requires double stranded DNA, particular to biological DNA replication.

**Figure SI 1.**
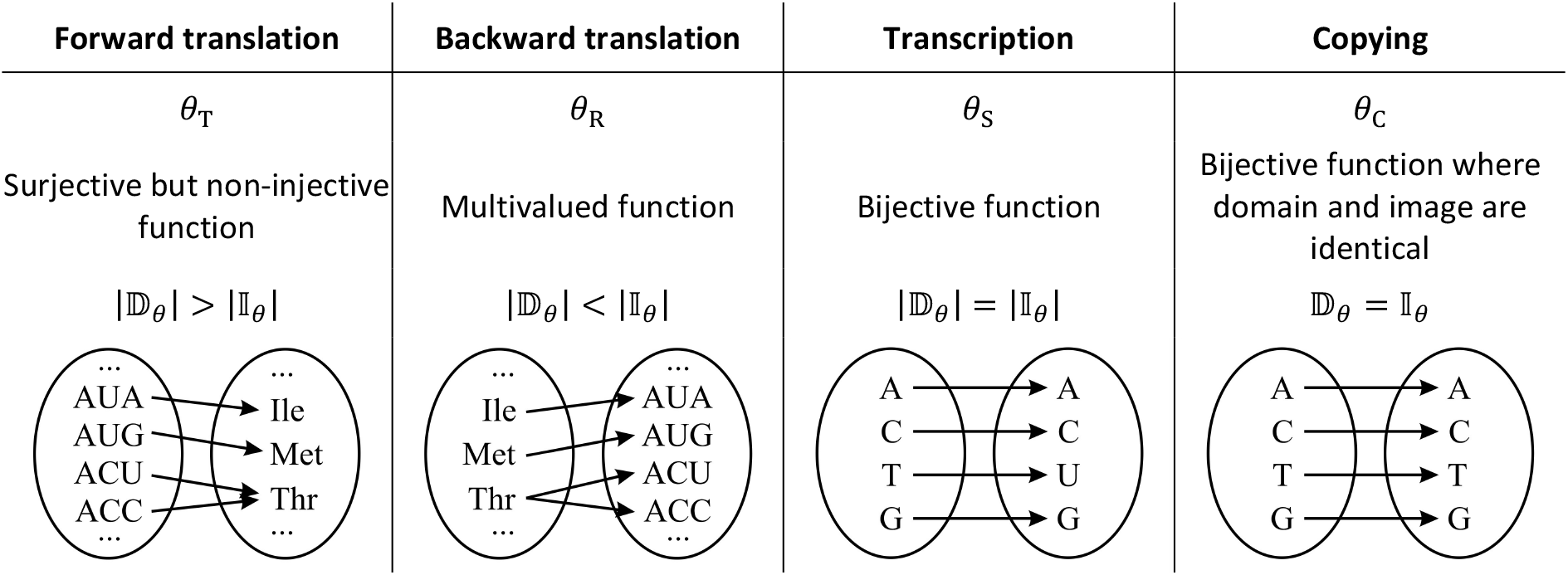
Elementary information transformations, their symbols, function descriptions, relationships of their domain and image, and examples. A transformation defines a unidirectional relationship between elements of the source set (domain) and elements of the target set (image). For sake of simplicity, we rely on terms related to polymer processing. Transformations can be applied to a list of elements (e.g., to codons) to be transformed (e.g., to a list of amino acids). The reverse translation of DNA is a hypothetic process.

If a transformation does not map an element of a domain to anything or maps multiple elements to the same target element, it loses information. We consider only transformations that map all possible elements of X to at least one element of Y, consistently (i.e., mapping does not change between successive transformations).

For example, 64 codons mapped to 20 amino acids by the standard genetic translation (an instance of *θ*_*t*_) is a reduction of information by ∼70 %. Thus, *θ* can maintain only as much information as is represented by the smaller set, domain 𝔻 _θ_ or image 𝕀 _θ_ (in this case, the 20 amino acids). That is, the potential maximal information maintainable by *θ* is *î* _θ_ = min(|𝔻 _θ_ |, |𝕀 _θ_ |).

Note, that the term “codon” is used here in a general way, as a module of a source polymer that directly maps to a module of the target polymer. For example, the codon space of DNA during translation consists of 64 nucleotide triplets (codons), while the corresponding codon-space of proteins contains 20 amino acids plus the stop signal. A function mapping the 64 codons to their 21 targets defines the elementary transformation *θ*_DNA→AA_, that can be used to transform a DNA sequence of any length (multiples of 3). In case of transcription, the codon spaces of source and target sets are of identical sizes (one containing DNAs the other RNAs). Therefore, an informational inheritance system can be represented by a transmission topology graph *G*, defined by the sets of states X and Y and the set of elementary transformations {*Θ*_l_, *Θ*_2_, … } defined between states (Figure 3 and Figure SI 3).

### Complex transmissions

To analyse complex information transmissions (those in Figures 2 and 3), we define the complex transmission Θ as the simplest general, indefinitely long, linear transformation chain of two distinct states *x*_i_ ∈ X and *y*_*j*_ ∈ Y as:

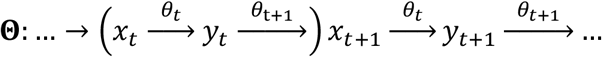

where *x*_*t*_ and *y*_*t*_ indicate the states at time step *t, θ*_*t*_ = *θ*_XY_ and *θ*_*t*+1_ = *θ*_YX_. Note, that the parenthesis denotes a repetitive subset of steps, modelled throughout this paper.

Θ may represent a replication process only if certain conditions are met. Here we assume that the successive *x*_*t*_ belong to the same medium, and the same is true for successive *y*_*t*_. Cases where *x*_*t*_ cannot be reconstructed at all are ignored, being not sufficient for replication. In case of exact template copying, one expects that *x*_l_ = *y*_l_ = *x*_2_ = *y*_2_ = …, i.e. all states are structurally identical. However, to maintain information against selection, the minimal condition is that **successive states are equivalent under selection**: *x*_l_ ∼ *y*_l_ ∼ *x*_2_ ∼ *y*_2_ ∼ …, i.e. phenotypes are identical under selection (for details, see ^4^). Accordingly, *v*(*x*_l_) = *w*(*y*_l_) = *v*(*x*_2_) = *w*(*y*_2_) = …, where *v* and *w* are fitness functions defined over sets X and Y, respectively. If state Y is under direct selection, generated from and representing *x*_*t*_, it must be true that *y*_*t*_ phenotypes are identical: *v*(*x*_*t*_) = *w*(*y*_*t*_), ∀*t*, hence the condition simplifies to *v*(*x*_*t*_) = *v*(*x*_*t*+1_), ∀*t*, that is, for an indefinite time. Relying on equivalency instead of identity allows other transformations than copying to be involved in the replication process. If *x*_*t*_ ≁ *x*_*t*+1_ and *y*_*t*_ ≁ *y*_*t*+1_, the transmission process is potentially losing information. If only one of the conditions is true, the system can only maintain as much information as the upholding condition allows (the equivalence on *x*_*t*_ *or* on *y*_*t*_).

Figure 4 describes potential complex information topologies corresponding the linear chain above, based on the generalization of Figure 3. In case of linear transmission chains, it is of no matter which state is autocatalytic, as ultimately, all states multiply. For simplicity, we assume that *x* is autocatalytic (ignoring surplus). Without further specification of the states *x* ∈ X and *y* ∈ Y, and ignoring mutation, the following distinct transmission topologies (i.e. pairs of successive transmissions) are identified:

- **Direct transmission**. All transformations are *copying, θ*_*t*_ = *θ*_u_ = *θ*_C_, ∀*t, u*, in which case *x*_*t*_ = *x*_*t*+1_ = *y*_*t*_ = *y*_*t*+1_, ∀*t*. Without mutation, the system cannot lose information.
- **Multistate transmission**. Successive transformations are *transcriptions*, and each one is the inverse of the previous one, 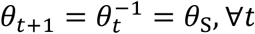, in which case *x*_*t*_ = *x*_*t*+1_ ⋀ *y*_*t*_ = *y*_*t*+1_, ∀*t*. Without mutation, the system cannot lose information. Topologically it is equivalent to direct replication.
- **Probabilistic transmission**. Successive transformations are not bijective but are (probabilistic) inverses of each other. That is, each transmission *θ*_i_ is a translation such that 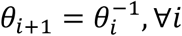 In other words, a *translation* is always followed by its appropriate *reverse translation*. There are two possibilities, identical up to a state shift (informationally isomorphic):
- 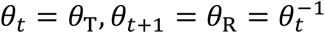, in which case *y*_*t*_ = *y*_*t*+1_, ∀*t*, but it is not necessarily true that *x*_*t*_ = *x*_*t*+1_. Thus, since *y*_*t*_ = *y*_*t*+1_, ∀*t*, the information content of state *y* can be stably maintained.
- 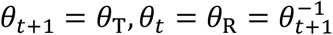, in which case *x*_*t*_ = *x*_*t*+1_, ∀*t*, but it is not necessarily true that *y*_*t*_ = *y*_*t*+1_. Thus, since *x*_*t*_ = *x*_*t*+1_, ∀*t*, the information content of state *x* can be stably maintained.

Since selection may act on one of the states only, there may be crucial differences (see later).

- **Information losing transmission**. Successive transformations are not (probabilistic) inverses of each other, 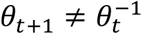, therefore, successive transformations cannot restore either state reliably. Consequently, there is no guarantee, that any state can be stably maintained, that is ∃*t, u*: *x*_*t*_ ≠ *x*_*t*+1_ ⋀ *y*_u_ ≠ *y*_u+1_. Therefore, the process inherently loses information, even if there is no mutation.

By dedicating *y* as the interactor, subject to selection, the following distinct transmission topologies are identified (Figure 4).

A. **Direct replication** (Figure 4A). *x* is a replicator with no intermediate state, hence no division of labour. Example is a (putative) prebiotic ribozyme, which is itself responsible for both conserving information and interfacing selection. The most advanced artificially selected polymerase ribozyme at the moment is of ∼200 nucleotides long and can only replicate sequences up to a length of 95, thus it cannot replicate itself ^136^. Examples include von Kiedrowski’s self-replicating oligomers, though with limited heredity ^137^.
B. **Replication with exact intermediate** (Figure 4B). Both *x* and its intermediate *y* can be maintained stably without loss of information. Therefore, both *x* and *y* are replicators, and *y* (subject to direct selection) is also an interactor. It lacks division of labour, as changes applied to *x* or *y* both can be inherited, thus there is no dedicated entity for either keeping information intact or interfacing selection. Any informational autocatalytic chemical cycle with at least one intermediate belongs here, for example replication of RNA +/- strands (where the + strand generates a - strand, that generates a + strand, and so on). If either *x* or *y* is omitted (as they do not convey extra information), the system simplifies to direct replication (Figure 4A).
C. **Weismannian replication** (Figure 4C). Division of labour between *x*, the dedicated replicator, and *y*, the dedicated interactor. Information introduced to *y* cannot propagate to *x*, and *y* alone is responsible directly for selection. The successive *y*_*t*_ need not be identical, but ideally their phenotypes are: *w*(*y*_*t*_) = *w*(*y*_*t*+1_), ∀*t* (when mutation is ignored). Example is the idealized gene-protein inheritance system.
D. **Successively Lamarckian replication** (Figure 4D). *y*_*t*_ is transformed to *x*_*t*+1_ ≠ *x*_*t*_, due to ambiguous processes, therefore it may not look like a replicator. However, since all *x*_*i*_ code for the same *y*, they are phenotypically equivalent, regardless of their actual structure. Thus, this is replication, but the only information that can be stably maintained is of *y* but not of *x*. There is only partial division of labour, as both *x* and *y* can inherit changes but only *y* is responsible for direct selection. A hypothetic example is the DNA → protein → DNA → … system, where a certain amino acid can be retranslated to DNA. Even if information is maintained, the evolution of such systems may fundamentally differ from the dynamics of direct or Weismannian replication.
E. **Probabilistic replication** (Figure 4E). *x*_*t*+1_ is translated to *y*_*t*+1_ ≠ *y*_*t*_. Since transformation of *x*_*t*_ is ambiguous, there is no guarantee that *y*_*t*+1_ (or any subsequent *y*_*t*+n_) translates to *x*_*t*_. Therefore, replication can only be stable, if two conditions are met. First, the phenotype of *x* must always be the same, that is *v*(*x*_*t*_) = *v*(*x*_*t*+1_) = *w*(*y*_*t*_) = *w*(*y*_*t*+1_), ∀*t*. Second, each *y*_*t*_ must yield the same genotype *x*. If neither condition is met, the process loses information (Figure 4F). If only the first condition is met, then whatever *x*_*t*+1_ the *y*_*t*_ is translated to, it is always true that *v*(*x*_*t*_) = *v*(*x*_u_), ∀*t, u*, hence only the information content of the phenotype can be maintained. If only the second condition is met, the process can replicate *x*, but with potentially different phenotypes every time. Presumably, evolution of such inheritance systems tends toward the more exact replication of Figure 4B, i.e., toward unambiguous translation if *x*.
F. **Information losing transmission** (Figure 4F). Neither state can be maintained stably, as due to informational loss, there is no guarantee, that *w*(*y*_*t*_) = *w*(*y*_*t*+1_) can be maintained for all *t*. Hence there is no stable replication. If, however, *w*(*y*_*t*_) = *w*(*y*_u_), ∀*t, u*, it simplifies to Probabilistic replication I. (Figure 4D). If all successive *y*_*t*_ are equivalent under selection, it also means that all successive *x*_*t*_ are equivalent too.

**Simultaneously Lamarckian transmission** (Figure 4G). Both states *x*_*t*_ and *y*_*t*_ contribute to the next generation *x*_*t*+1_, according to some unspecified rule deciding how individual codons (or elements) contribute in parallel (or not at all). Again, depending on the relation of *θ*_l_ and *θ*_2_, replication can be more or less exact (as in case of Figure 4F). We have modelled such a transmission system in Appendix 6 (also see Figure 5).

While codon-based informational compositionality seems to be specific to polymer-based inheritance, we can safely assume that cultural transmission can also be represented as a chain of elementary transformations, based on two reasons. First, holistic transformation is the natural extremum of compositional transformation, when *n* = 1 (and the codon space is as big as the sequence space). Second, parts of culture are clearly compositional in nature, most notably lexicon and grammar. Early language might have been less compositional as it is today (cf. holistic protolanguage ^138–140^), but for a baseline assumption, we maintain that parts of cultural inheritance can be represented by compositional transformations as modelled in this paper.

### Examples of complex transmissions

The elementary transformation of forward translation *θ*_*t*_ can be specified in various ways. A uniformly ordered *θ*_*t*_maps the ordered set of source codons to the ordered set of target codons with uniform probability, evenly distributing target codons among source codons (Table SI 2A). Uniform random *θ*_*t*_ distributes target codons among source codons evenly, though not in successive order (Table SI 2B). These two are qualitatively identical, up to a re-indexing of source and/or target codons. Random *θ*_*t*_ does not necessarily distribute evenly (Table SI 2C).

**Table SI 2.** Possible forward translation methods of the elementary transformation θ_XY_, that map the codon space S_X_ of size 8 to the codon space S_Y_ of size 2 (c_X_ = 3, c_Y_ = 1, a_X_ = a_Y_ = 2. **A**: uniformly ordered θ; **B**; uniformly random θ; **C**: random θ.

| **A**<br>$\theta_{\text{uniformly ordered}}$ | **B**<br>$\theta_{\text{uniformly random}}$ | **C**<br>$\theta_{\text{random}}$ |
| --- | --- | --- |
| 000 → 0 | 000 → 0 | 000 → 1 |
| 001 → 0 | 001 → 1 | 001 → 0 |
| 010 → 0 | 010 → 1 | 010 → 1 |
| 011 → 0 | 011 → 0 | 011 → 1 |
| 100 → 1 | 100 → 0 | 100 → 0 |
| 101 → 1 | 101 → 1 | 101 → 1 |
| 110 → 1 | 110 → 0 | 110 → 1 |
| 111 → 1 | 111 → 1 | 111 → 1 |

An appropriate inverted backward translation 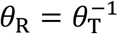 must be probabilistic in each of the examples to maintain maximum information. Throughout this paper, simple probabilistic choice is used to resolve backward translation: one of the possible target codons is chosen randomly, each with equal probability. Such *θ*_R_ maps the same Y codon (0 or 1) into multiple X codons with equal probability (e.g., in Figure SI 2C, the Y element Valine maps into multiple X codons (GUU, GUC, GUA, GUG)).

**Figure SI 2.**
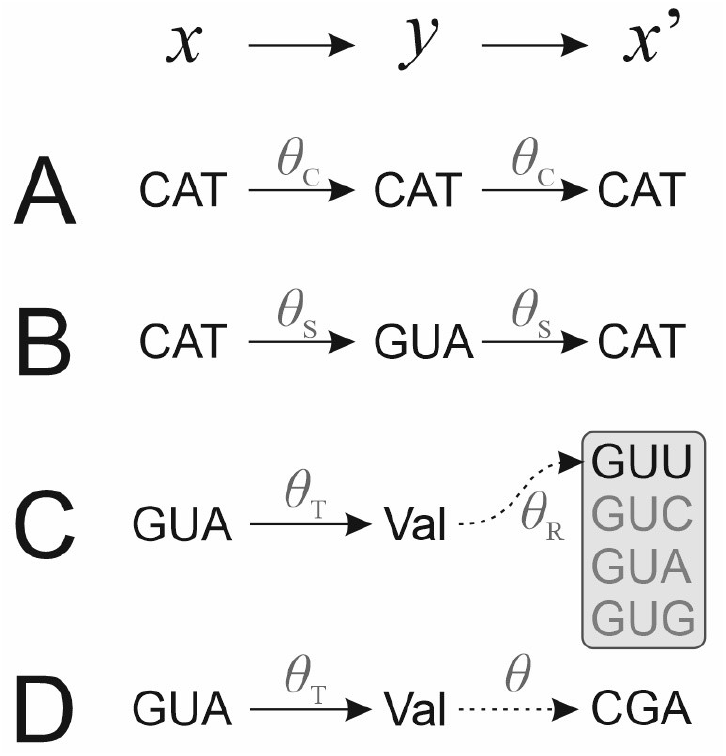
Different realizations of the Lamarckian topology x → y → x’ of Figure 3C assuming different θ_XY_ and θ_YX_ transformation processes. **A**: Consecutive transformations are all copying that do not change the inventory of the medium. **B**: Consecutive transformations are transcriptions that change the medium but not the inventory size. **C**: Consecutive transformations are probabilistic inverses of each other: the information of one state can be maintained, as all codons code for valine. **D**: Consecutive transformations are not probabilistic inverses of each other, hence information cannot be maintained, as CGA does not code for valine.

We also investigate the case where the successor of a forward translation is not its probabilistic inverse, i.e.,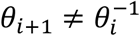. In such cases information is inherently lost in successive transformations. Figure SI 3 shows some example cases, where the backward translation is the inverse of the forward translation (columns 1, 2) or not (columns 3, 4). Information can only be maintained in successive pairs of transformations without loss if the backward translation is the inverse of the forward translation. In the other case, there may be partial inversion that may ensure the stable inheritance of parts of codon spaces. There may be chance that more than one successive translation-retranslation pair can return a codon to its own equivalency class, but that means that there are actually less phenotype classes than expected. At the end of the continuum is a disjunct pair of *θ*_i_ and *θ*_i+1_ where no codon can be translated back to its own equivalency class.

**Figure SI 3.**
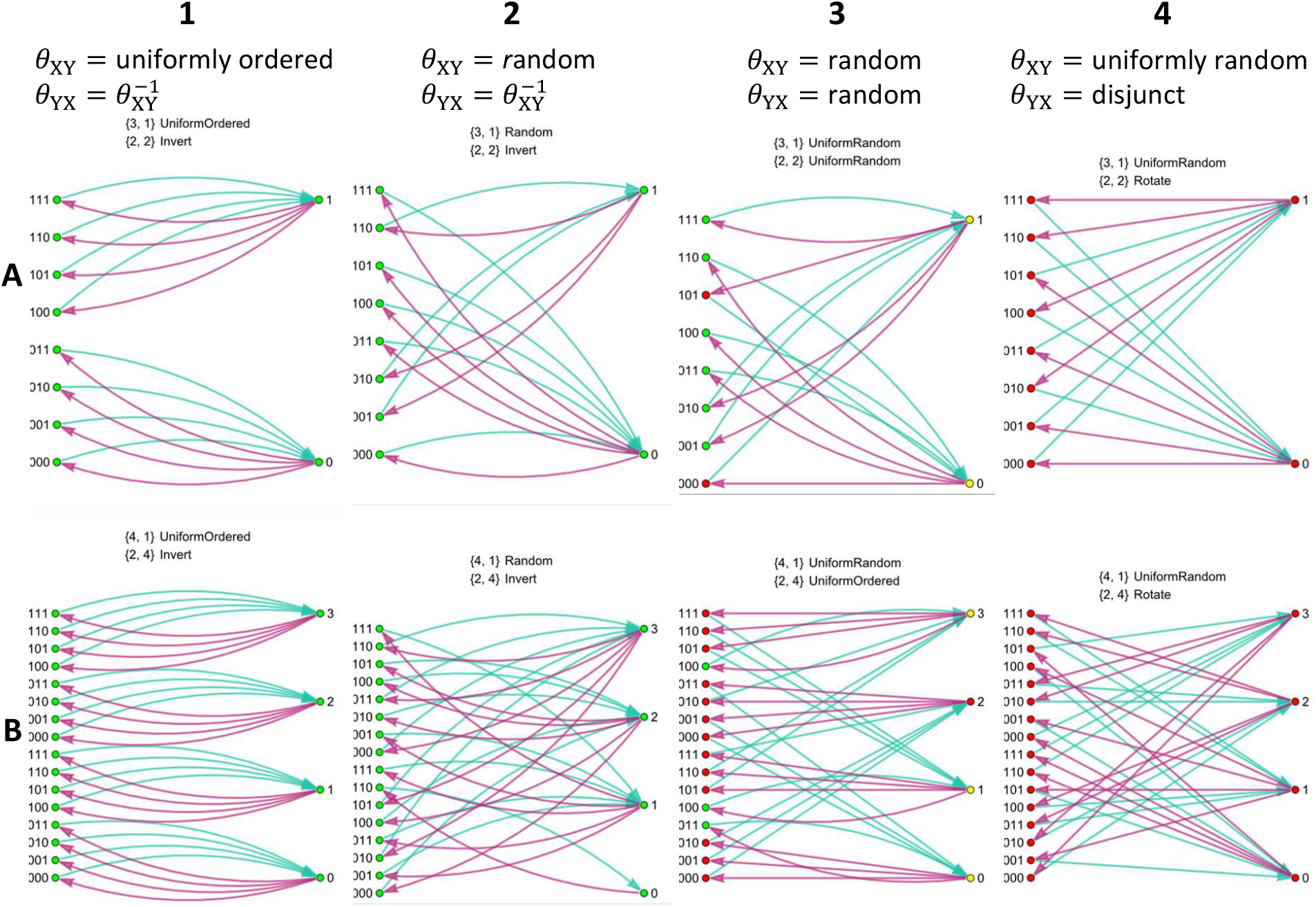
Relation of successive forward and backward transformations. On each graph, the left column lists elements x_i_ ∈ S_X_, right column the elements y_i_ ∈ S_Y_. Parameters are **A**: {c_X_ = 3, c_Y_ = 1, a_X_ = a_Y_ = 2}; **B**: {c_X_ = 4, c_Y_ = 1, a_X_ = 2, a_Y_ = 4}. Nodes are colored green, if the corresponding element x_i_ maps to a target y_i_ that maps back to x_j_, x, … that are all phenotypically equivalent to x_i_ (that is: x_i_ ∼y_i_ ∼x_j_ ∼x ∼ …); yellow, if the target y_i_ maps out of the original phenotype class of x_i_; and red if an element is not mapped back to its own phenotype class at all.

Instead of focusing on the actual per-digit copying fidelity in replication of Weismannian and Lamarckian replicators, one must realize that it is the Y state, the state under selection, that is ultimately important from an evolutionary point of view, and should be tracked during replication (according to the **principle of phenotypic equivalency** ^4^). Now the Y state might consist of less digits than the X state, due to the nature of translations. This renders calculations needlessly cumbersome. However, both X and Y consist of the same number of codons. Accordingly, we must deal with the copying fidelity of *codons* instead of single *digits*. For analysis, this simple inheritance system is parameterized according to Table SI 3.

**Table SI 3.**
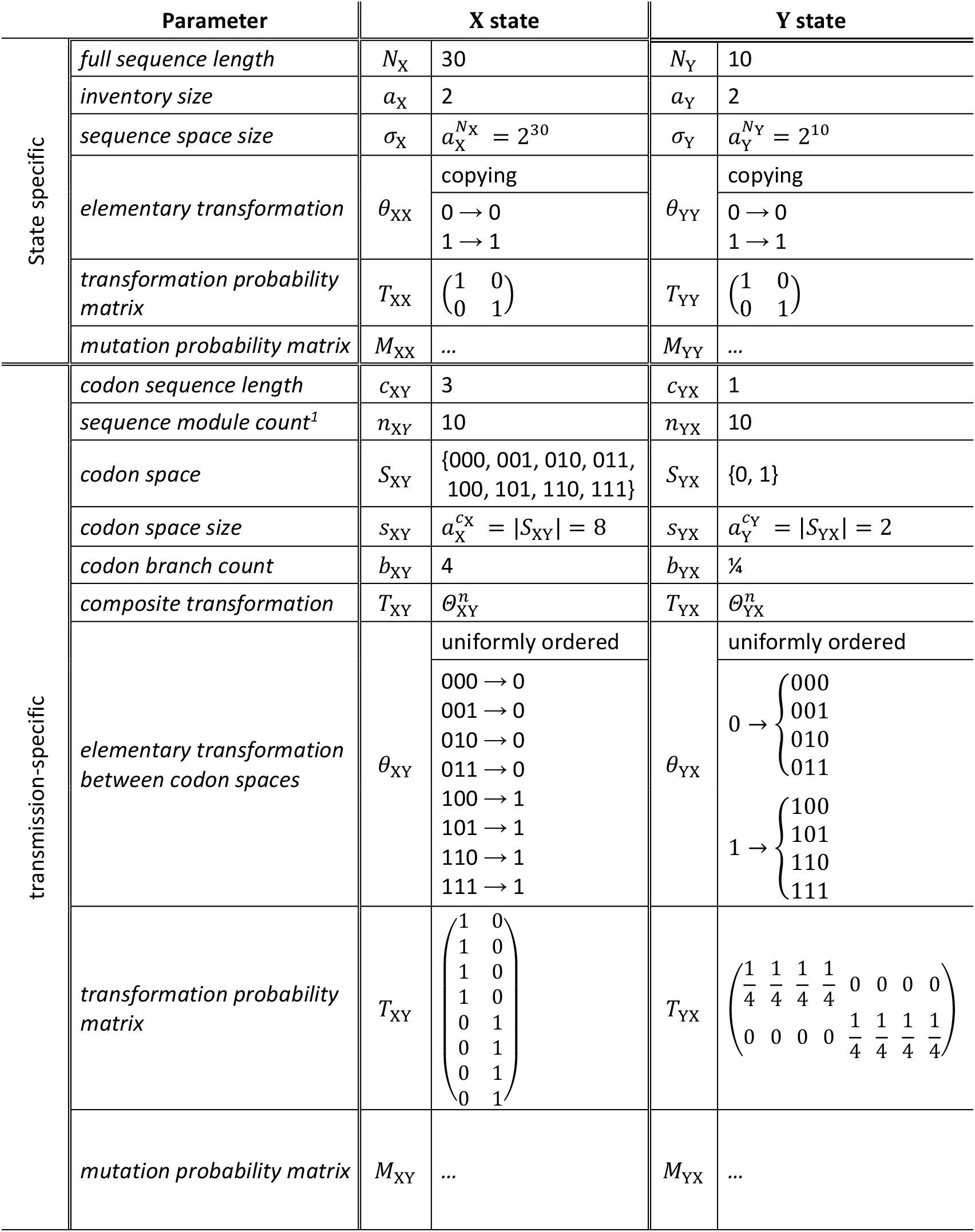
Parameters and properties of an example two-state (X, Y) inheritance system. For various θ, see Table SI 2 and Figure SI 3A, for T and M, see Figure SI 5. For replication probability surface, see Figure SI 7C. ^1^ The sequence module count is uniform for all transmissions: n = n_XY_ = n_YX_.

As Lamarckian inheritance has the Y sequence as the stable state during successive replication events, it is necessary to compare the overall replication probability of the Y sequences to the replication probability of the Y sequence during Weismannian replication. Furthermore, sequence replication fidelity depends on the codon replication probability and the number of codons in a sequence. Therefore, it is sensible to simplify the case to measure codon replication fidelity on the first hand. Using the model, I intend to answer the question: *What is the probability P*_*r*_ *that replicating an* X *codon yields the same* Y *codon, as is the original* Y *codon of the original* X *sequence?*

To understand the difference between the two topologies, each informational route is broken down to elementary transformations, where mutation always acts on the result of a transformation, as is depicted in Figure SI 4.

**Figure SI 4.**
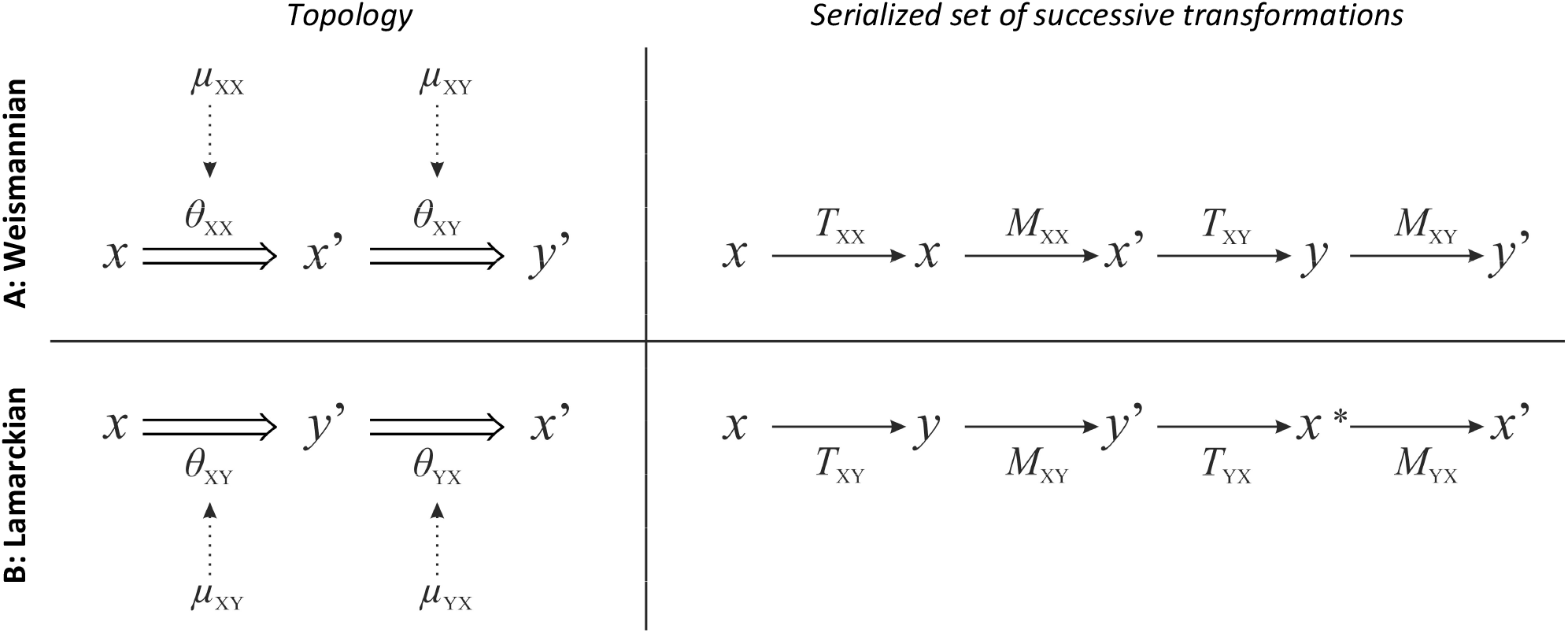
Detailed informational topologies, generating y ∈ Y from x ∈ X. **A**: Weismannian inheritance. **B**: Lamarckian inheritance. Left column: linearized version of Figure 3. Right column: Order of application of transformation (T) and mutation (M) matrices. θ_ij_ denotes the elementary transformation from state i to state j, T_ij_ denotes the transformation probability matrix of the codon space i to j corresponding to θ_ij_, μ_ij_ denotes the per-digit mutation rate applied to state j during transformation i → j, M_ij_ denotes the mutational probability matrix of the codon space S_j_, dependent of mutation rate μ_ij_.

## 3. Probability of Weismannian replication

To calculate the probability value, we introduce some matrices. Let the transformation probability matrix *T*_ij_ represent the transformation *θ*_ij_ from state *i* to state *j* as a matrix of size *s*_i_ × *s*_*j*_ of which the element *T*_ij(pq)_ gives the probability of transforming the *p*^th^ codon of the codon space of state *i* to the *q*^th^ codon of the codon space of state *j*. For translation (or transcription or copying), this matrix has 1 and 0 as its elements, while for reverse translation 0 and 1/*b*, where *b* is the codon branch value such that *b*∈ℕ and *b* ≥ 1. For irregular translations, *b* is defined as the sum of appropriate columns of *T*_ij_. We define the elementary process of copying as the identity function, hence *T*_ii_ is the identity matrix of size *s*_i_.

The codon mutation matrix *M*_*j*i_ is a square matrix of size *s*_i_ × *s*_i_ of which the element *M*_*j*i(pq)_ defines the probability that the *p*^th^ codon of the codon space of *i* is mutated to the *q*^th^ codon of the same space, assuming a per-digit mutation rate of *μ*_*j*i_. More precisely, the probability of mutating sequence (or codon) *p* to sequence (or codon) *q* is:

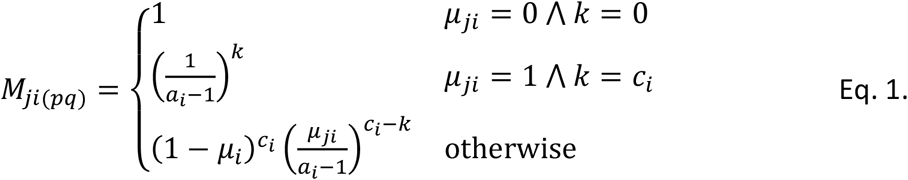

where alphabet size *a*_i_, codon length *c*_i_ and mutation rate *μ*_*j*i_ are specific for the *i*^th^ state, and *k* is the Hamming distance of sequences *p* and *q*. Examples of *T* and *M* matrices for the parameter configuration of Table SI 3 are shown in Figure SI 5.

**Figure SI 5.**
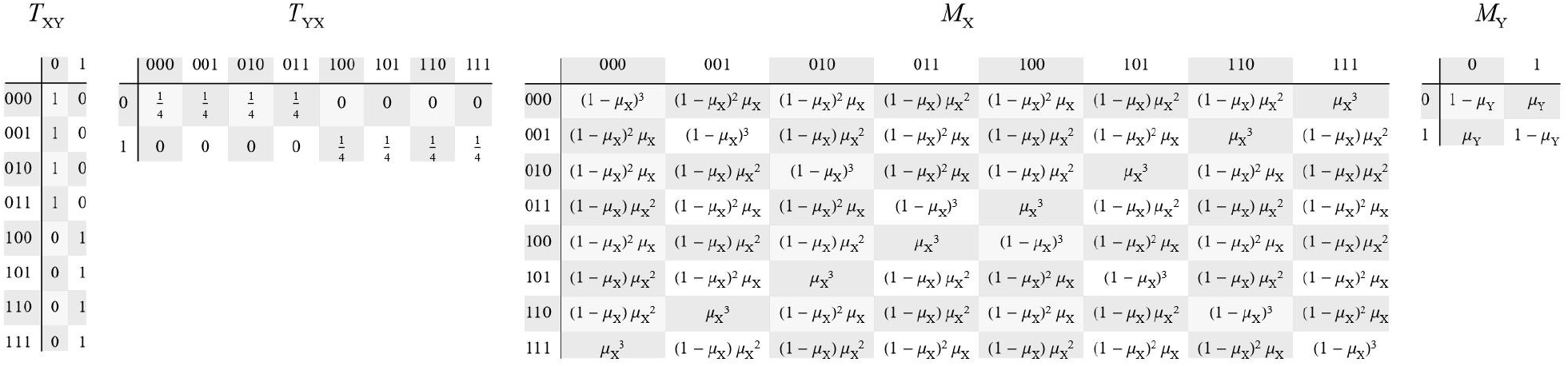
Examples of transformation probability matrix T and mutation probability matrix M. Cell at i^th^ row and j^th^ column gives the probability that the codon of the i^th^ row is transformed or mutated to the codon of the j^th^ column. Parameters are {c_X_ = 3, c_Y_ = 1, a_X_ = a_Y_ = 2}. The mutation matrix {M_X_, M_Y_} may stand for {M_XX_, M_XY_} in case of Weismannian replication or for {M_YX_, M_XY_} for Lamarckian replication.

First, we formalize codon replication probability *P*_r_ for the Weismannian topology. Let *P*_W_ be the probability that an *x*_i_ ∈ *S*_X_ codon is replicated such, that it returns the same *y*_*j*_ ∈ *S*_Y_ codon the original was coding for. First, let us define the **codon transformation probability** *ω*_ij_ that the *i*^th^ codon *x*_i_ is copied, mutated, translated, and again mutated to yield the *j*^th^ codon *y*_*j*_:

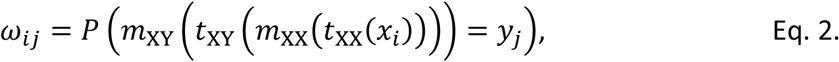

where *t*_ij_ and *m*_ij_ are the functional forms of *T*_ij_ and *M*_ij_, respectively. We have defined *t*_XX_ as the identity function for copying, hence it can be ignored (but not the mutation function *m*_XX_ of copying). Since *ω*_ij_ must be defined for all codons of *S*_X_ and *S*_Y_, it is convenient to write *ω* of size *s*_X_ × *s*_Y_ in matrix notation:

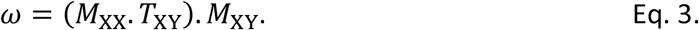

The **average codon replication probability**, *P*_W_, can be calculated by multiplying the codon transformation probability matrix *ω* with *T*_XY_ (this factors out the impossible translations), and averaging the result over all codons of *S*_X_. *P*_W_ gives the average probability that an arbitrary *x*_i_ codon is replicated faithfully to yield an *x* codon phenotypically equivalent to *x*_i_. That is, *both x*_i_ *and x encode the same y*_*j*_:

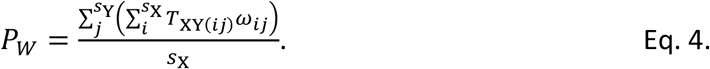

### 4. Probability of Lamarckian replication

Similarly to *ω*_ij_ for Weismannian replication, we define the **codon transformation probability** *λ*_ij_ for Lamarckian replication, following the Lamarckian topology of information transmission. *λ*_ij_ defines the probability of codon *x*_i_ being translated, mutated, retranslated, and again mutated to yield codon *x*_*j*_:

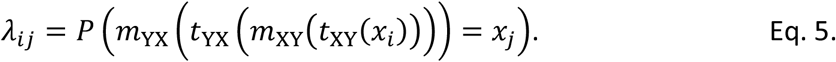

As there is no direct copying involved, no elementary transformation can be ignored. Also, the mutation functions are different compared to the Weismannian case before: *m*_XY_ and *m*_YX_, according to Figure 3 and Figure SI 4. The resulting matrix notation is:

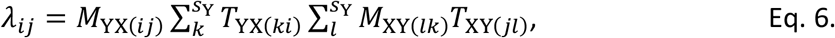

from which the **average codon replication probability** is:

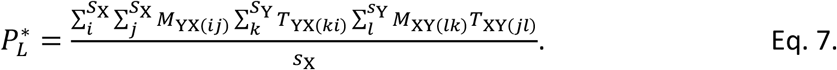

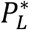 gives the probability of exactly replicating *x*_i_. However, since the Lamarckian state can only maintain as much information as is stored in the Y state, we are interested in the replication of the *y* ∈ Y state. Thus, we seek the *probability that the outcome of replication is phenotypically equivalent to the source*, i.e., when *x*_*j*_ stays in the same neutral subspace where the parent *x*_i_ was. In other words, what is the probability that *x*_*j*_ translates to the same thing *x*_i_ translates to (ignoring mutation)? The **average phenotypically equivalent codon replication probability** is:

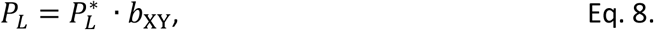

where *b*_XY_ is the average number of codons in *S*_X_ assigned to a codon in *S*_Y_ (codon branch value). This is an approximation, assuming that codon assignment is evenly distributed (i.e., the same amount of source codons translates to a target codon and vice versa), which might not be the case in reality.

Codon robustness (defined by codon branch value *b*_XY_) naturally translates to sequence robustness, *β*_XY_, which is the number of sequences allocated to the same target sequence in regard of the given transformation. It can be calculated as:

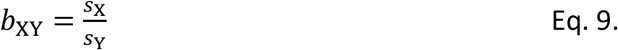

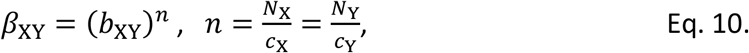

where *n* is the codon count of sequences (module count), that is identical for both X and Y sequences in any deterministic transformation model.

### 5. Probabilities of accurate replication

Probabilities of accurate replication were tested against iterations of sequences undergoing transformation and mutation, according to the following setup. For codons of the Weismannian topology, we have iterated over the range (0, 1) for *μ*_XX_ and *μ*_XY_, and for each {*μ*_XX_, *μ*_XY_} pair 1000 random codons were chosen from the codon space *S*_X_, and for each codon *x*_i_ ∈ *S*_X_ of the sample, the following equality was checked:

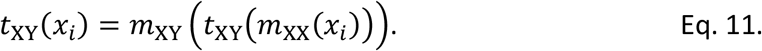

An example of the agreement between calculated and measured probability values of the Weismannian topology is illustrated in Figure SI 6A.

For codons of the Lamarckian topology, we have iterated over the range (0, 1) for *μ*_YX_ and

*μ*_XY_. For each {*μ*_YX_, *μ*_XY_} pair, 1000 random codons were chosen from the codon space *S*_X_, and for each codon *x*_i_ ∈ *S*_X_ of the sample, the following equality was checked:

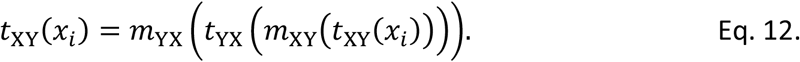

An example of the agreement between calculated and measured probability values of the Lamarckian and Weismannian topologies, and the agreement between these two topologies in case of lossless uniformly ordered transformations 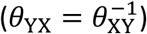 is illustrated in Figure SI 6B. For no mutation, one can observe what is naively expected: inheritance is exact, and perfect. With mutation, results show that for almost all cases Weismannian and Lamarckian probability surfaces have a near perfect agreement (Figure SI 7). For random but inverted transformations the difference becomes slightly pronounced but is still negligible Figure SI 7B, D.

**Figure SI 6.**
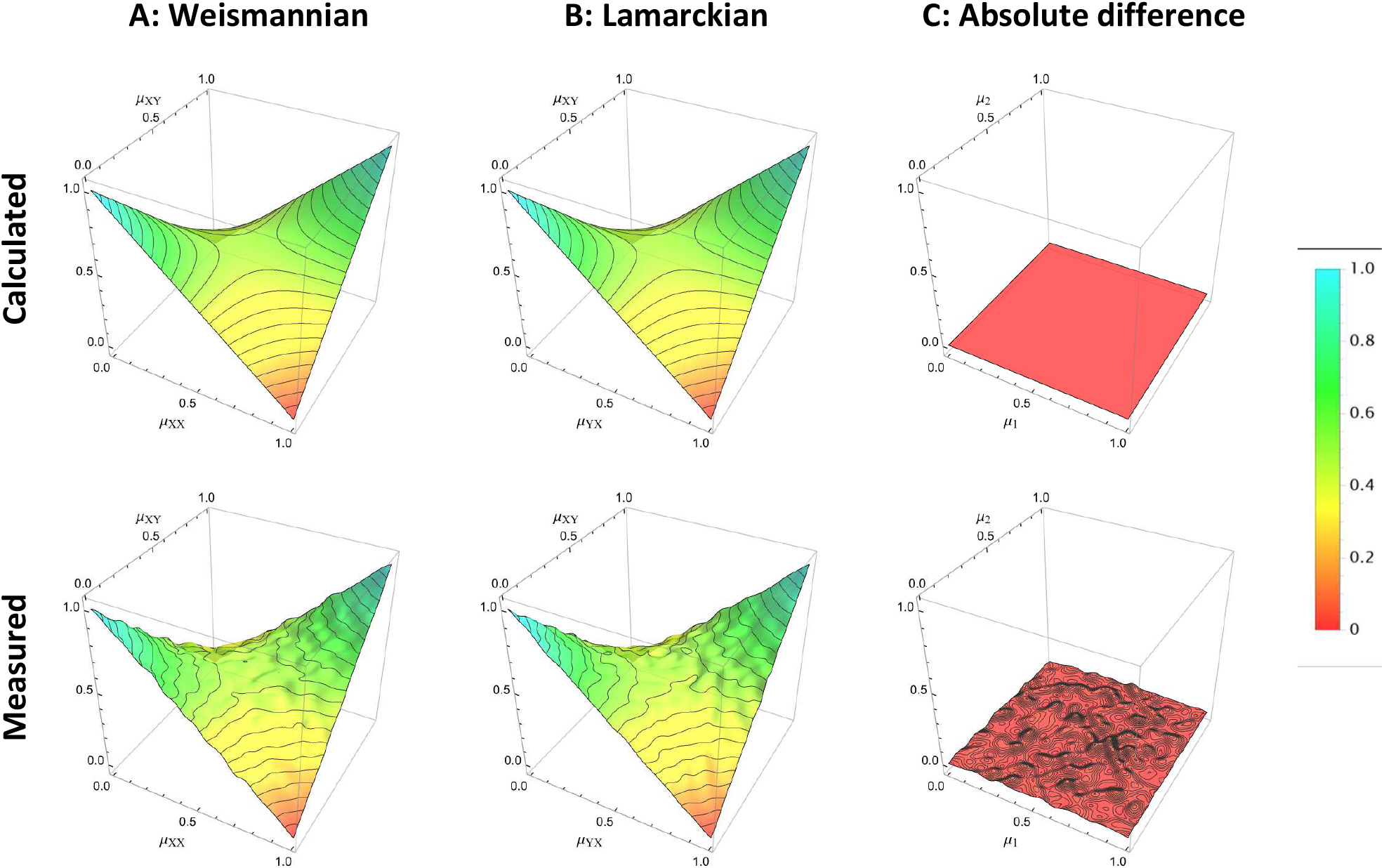
Average codon replication probability for Weismannian and Lamarckian inheritance, depending on the per-digit mutation rates of successive transformations in case of inverted backward transformation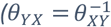, i.e., without transmissional loss of information. **A**: Weismannian replication probability P_W_ (according to Figure 3A, {μ_l_, μ_2_} = {μ_XX_, μ_XY_}); **B**: Lamarckian replication probability P_L_ (according to Figure 3B; {μ_l_, μ_2_} = {μ_YX_, μ_XY_}). The difference is negligible due to the uniformly ordered nature of translation θ_XY_ and the lossless inverted backward transformation of the Lamarckian topology. Top row: analytically calculated probability values according to Eqs. 4 & 8. Bottom row: measured probability values; for each value of μ_l_ and μ_2_, 1000 random iterations were performed to measure the probability. Both topologies rest on the same transformations and parameters: codon length c_X_ = 3, c_Y_ = 1, a_X_ = a_Y_ = 2, θ_XY_ = θ_uniformly ordered_,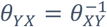.

The equality of probability surfaces breaks down the moment the second translation *θ*_YX_ is not the inverse of the first translation *θ*_XY_ (Figure SI 8). This has catastrophic effects for the Lamarckian topology where that the two successive transformations do not constitute an information-maintaining process anymore. Replication becomes inherently lossy, independent of mutation rates. In such cases, the replication probability of a Lamarckian replicator (for almost all parameter combinations) is below 1. This shows that inherently information-losing transformations cannot perform exact copying stably for all sequences **even if there is no exogenous mutation rate** during inheritance. Therefore, the two sources for information loss, external *mutation* and internal *transformational degradation* must be distinguished.

**Figure SI 7.**
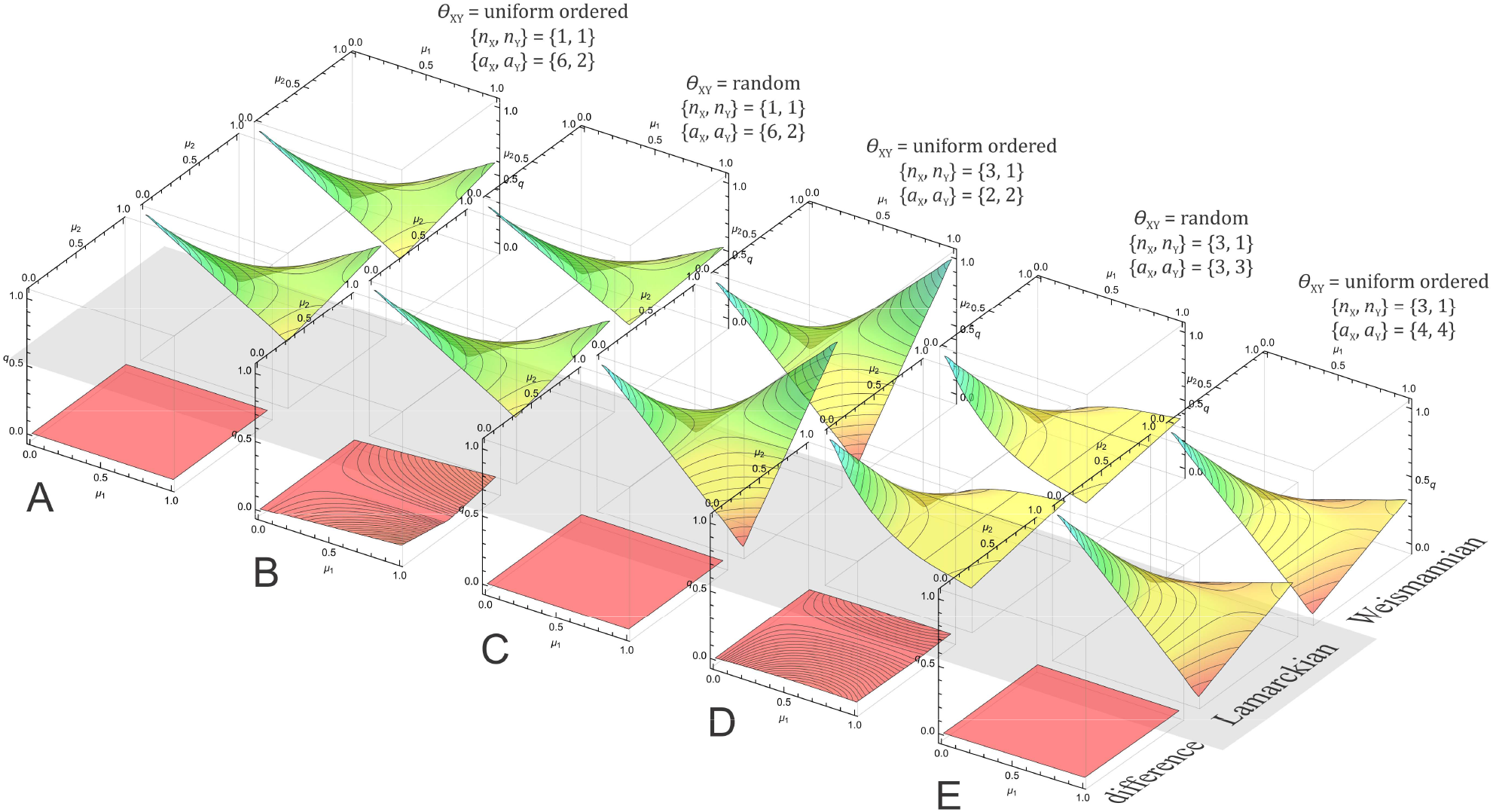
Average codon replication probability for lossless transmissions, measuring the probability that after replication, y_t+1_ phenotypically equals to y_t_. For each parameter set, the three plots are: Weismannian P_W_, Lamarckian P_L_, and their absolute difference: |P_W_ − P_L_ |. For each forward transformation θ_XY_, the backward transformation θ_YX_ is defined so that 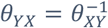. Note, that in case of Weismannian replication, there is no θ_YX_ involved. For P_W_ : {μ_l_, μ_2_} = {μ_XX_, μ_XY_}, while for P_L_ : {μ_l_, μ_2_} = {μ_YX_, μ_XY_}, that is, the axes for μ_XY_ are the same.

**Figure SI 8.**
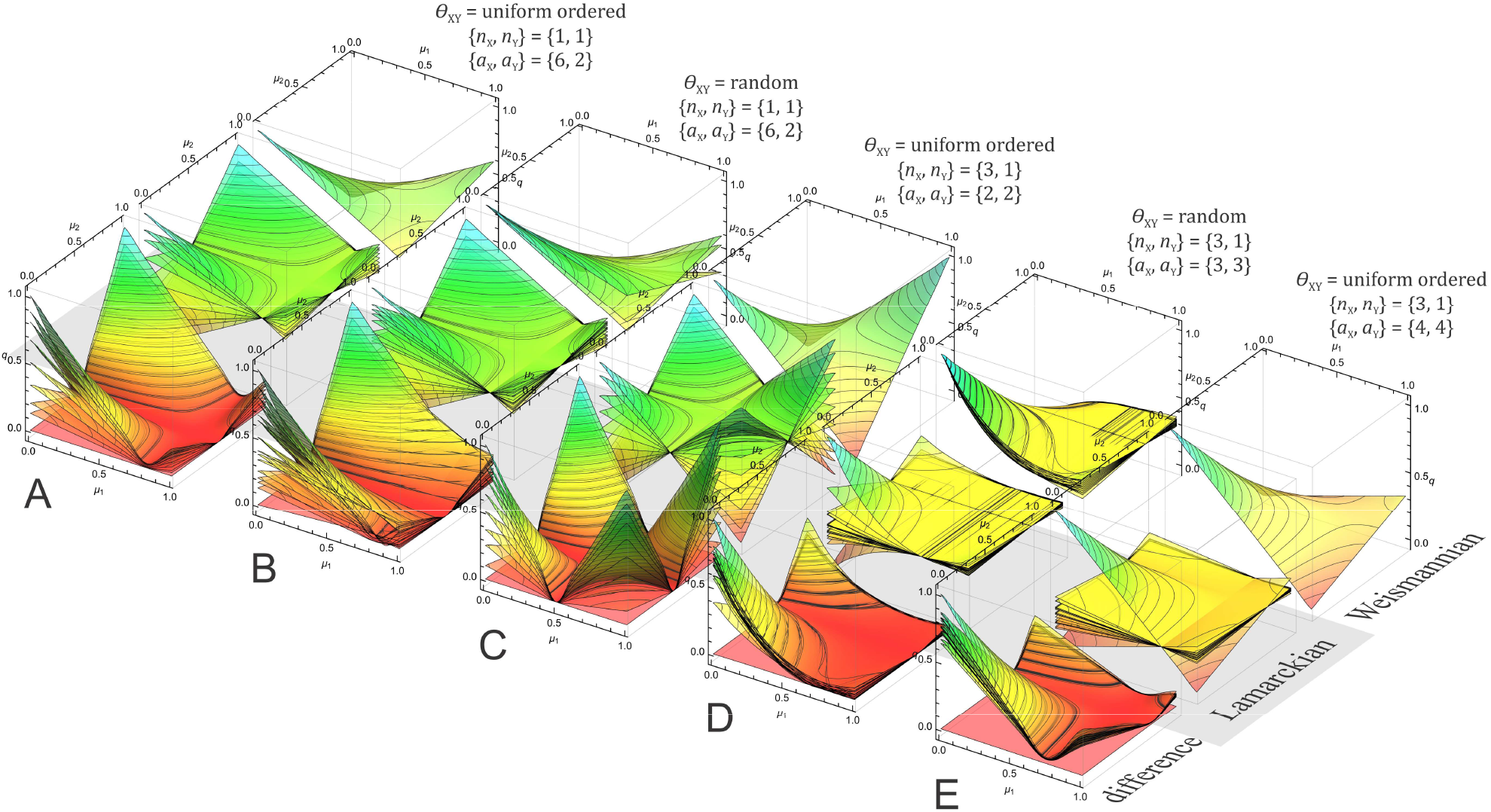
Average codon replication probability for information-losing transmissions, measuring the probability that after replication, y_t+1_ phenotypically equals to y_t_. For each parameter set, the three plots are: Weismannian P_W_, Lamarckian P_L_, and their absolute difference: |P_W_ − P_L_ |. For all plots, 20 independent forward transformations θ_XY_ were generated, and for each θ_XY_ a random backward transformation θ_YX_ was generated so that 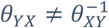. The replication probabilities corresponding to each pair of {θ_XY_, θ_YX_} are shown as multiple surfaces within the same box. Note, that in case of Weismannian replication, there is no θ_YX_ involved and a uniformly ordered forward translation θ_XY_ is deterministically defined, hence the multiple surfaces do not differ much. For P_W_ : {μ_l_, μ_2_} = {μ_XX_, μ_XY_}, while for P_L_ : {μ_l_, μ_2_} = {μ_YX_, μ_XY_}, that is, the axes for μ_XY_ are the same.

For small inventory sizes, there is more chance to achieve tolerable overall error rates, as there is a higher chance that two consecutive mutations bring the given bit back to its original state. As an extremum, at *a* = 2 (i.e. two-letter alphabet), there are two cases for which exact copying can be reached. Furthermore, only for *a* = 2 is it possible to have 0 possibility of exact replication: when only one of the two transformations is erroneous but not the other, mutation inevitable destroys information (see Figure SI 7, front row, leftmost plots).

Thus, in case of the Lamarckian topology, smaller inventory size allows for larger replication fidelity, as with increasing alphabet size overall fidelity drops to a general level, mostly independent of the mutational rates at all (Figure SI 8, rightmost plots). However, probabilities of Lamarckian topology could never get close to the probability surface of the Weismannian topology in such lossy complex transmissions. The only chance for it is if the reverse transformation table is – by chance – randomly generated to be the inverse of the forward transformation table. This case would then be equal to the *uniform random* method of Table SI 2.

### 6. Population-based stochastic simulation

Stochastic simulations support the above results. A simple stochastic replication-selection system was built to check the difference of Weismannian and Lamarckian inheritance. We have followed the most general topology (see Figure SI 9), where either *x, y*, or a combination of them defines the next generation of *x* (*y* is always generated form *x*). Accordingly, we have introduced parameter *α* ∈ (0,1) that defines the ratio of *x*_*t*_ and *y*_*t*_ in defining *x*_*t*+1_: if *α* = 1, it is solely *x* that generates *x*_*t*+1_; if *α* = 1, only *y*_*t*_ generates *x*_*t*+1_; any intermediate *α* means that the first *α* part (rounded) of *x*_*t*_ and the last (1 − *α*) part (rounded) of *y*_*t*_ defines *x*_*t*+1_. For sake of simplicity, all sequences are assumed to be binary (*a*_X_ = *a*_Y_ = 2). Sequence lengths were chosen to fit both transformational requirements (i.e., a large-enough codon branching value was chosen: *b*_XY_ = 3), and the population size, so we see the effect of reduced X-diversity when switching from Lamarckian to Weismannian replication. Mutation rates were chosen to yield identical sequence-replication chances for Y, according to previous results: *μ*_XX_ = *μ*_XY_ = *μ*_YX_ = 0.01.

A standard selective scenario was used, where the population settles at the best fitness due to selective birth (replication) and random death. Fitness is defined by the combination of subfitness values *w*_X_ and *w*_Y_ calculated for the X and Y sequences, respectively, as the relative Hamming distance compared to the arbitrary best sequence defined for X and Y. We have selected opposing best sequences: all-1 for all *x*_master_ ∈ X and all-0 for y_master_ ∈ Y. The fitness of a member of the population is the linear combination of the subfitness values, defined by the mixing parameter *β*:

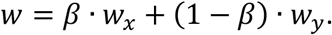

At each replication event, a member of the population (defined as the pair {*x*_*t*_, *y*_*t*_}) is randomly chosen, with probability proportional to its fitness *w*. {*x*_*t*_, *y*_*t*_} is then replicated to yield the new *x*_*t*+1_, according to the mixing parameter *α* that defines in what amount *x*_*t*_ and *y*_*t*_ contribute to *x*_*t*+1_ (and mutation rates *μ*_XX_ and *μ*_YX_). For sake of simplicity, we copy the first *α* · *n* · *c*_X_ elements (rounded) of *x*_*t*_ and reverse translate the last (1 − *α*) · *n* · *c*_Y_ elements (rounded) of *y*_*t*_, and joins them to form *y*_*t*+1_. The new *y*_*t*+1_ is then created from *x*_*t*+1_, with mutation rate *μ*_XY_. The new replicant pair {*x*_*t*+1_, *y*_*t*+1_} then randomly replaces a member of the population.

**Figure SI 9.**
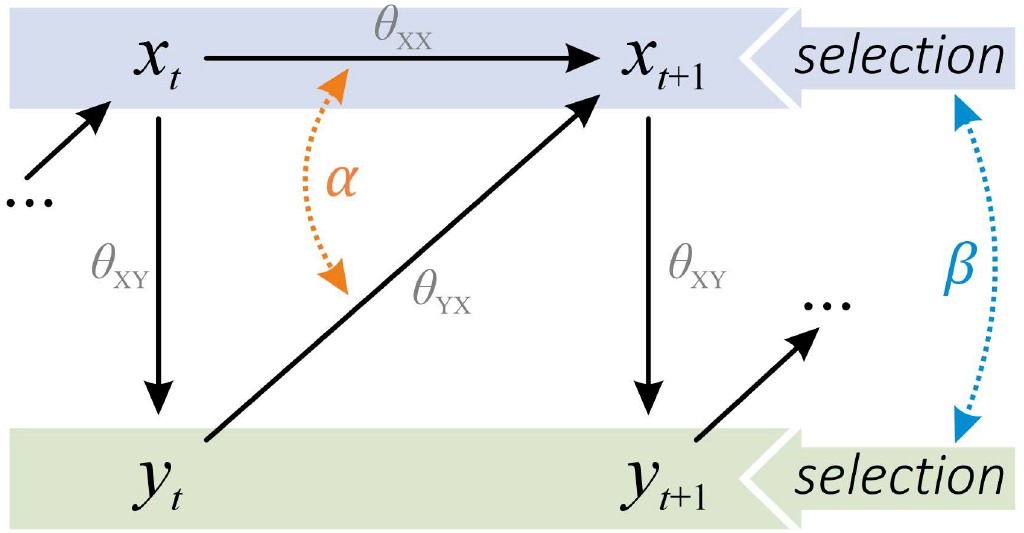
General replication topology of two information states x ∈ X and y ∈ Y, following Figure 4G of the main text. The next generation of x_t+1_ is generated by combining the copied and reverse-translated information contents of x_t_ and y_t_ according to the mixing parameter α. Selection is defined by the fitness of {x_t_, y_t_} pairs as the linear combination of the theoretical subfitness values w_x_ and w_y_, defined by the mixing parameter β as w = β w_x_ + (1 − β) w_y_.

Results indicate that populations reach optimal fitness with roughly the same pace and the equilibrium diversities of Y-sequences are identical (Figure SI 10-11, also see Figure 5). It cannot settle at 1, as the mutation rate guarantees the continuous emergence of new variants, but it gets very close to it. The only distinct difference between the two simulations is in the magnitude of diversity of X sequences. Without mutation, diversity *d*_x_ (the number of different X sequences in a given timestep) approximates sequence robustness 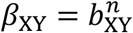 both for Weismannian and Lamarckian inheritance (see Eq. 10, not shown here), which quickly outgrows the actual population size (for this particular case, *β*_XY_ = 1024). A smaller population size reduces actual *d*_X_ as it limits *d*_Y_ of the population, while mutation rate increases it, hence it is not trivial how to calculate the actual equilibrium diversity in a given case.

**Figure SI 10.**
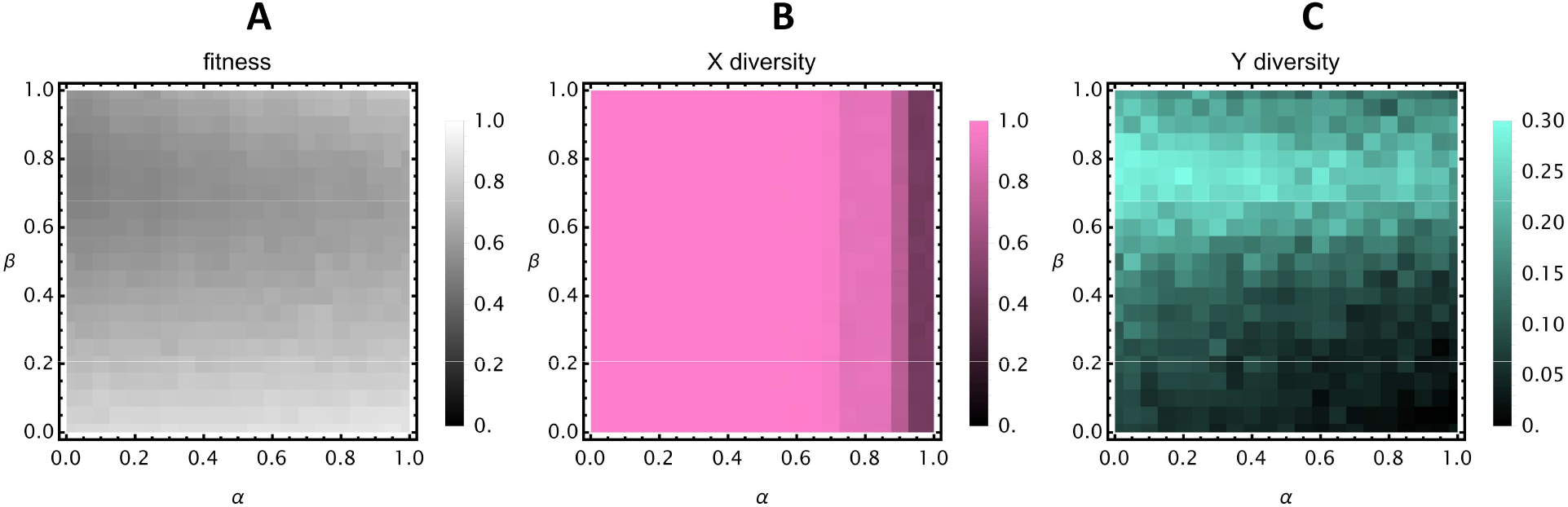
Average fitness and X- and Y-diversities of the Lamarckian-Weismannian continuum depending on inheritance-mixing (α) and interaction-mixing (β) parameters. Each pixel is the average of the equilibrium of a simulation with a specific {α, β} pair. **A**: The fitness of the population is always close to the maximum. **B**: X-diversity decreases as the replication topology gets closer to Weismannian replication. **C**: Y-diversity increases in Lamarckian populations: the more Y contributes to selection, the more diverse the Y-population is. Population size is 1000, diversity is scaled from (0,1000) to (0,1). For details and parameters, see Figure SI 11.

These stochastic simulations confirm that a population of Lamarckian replicators, even with fully variable intermediate X sequences, could maintain relevant information and evolve just like a population of Weismannian replicators evolve (Figure SI 11, e.g. *β* = 1.0, *α* = 0.5). The difference between the Weismannian and Lamarckian inheritance in this setup is purely in the amount of robustness of the system: the X-sequence diversity is always larger for the Lamarckian replicator due to the underlying larger codon robustness. To maintain the same amount of Y-information by Lamarckian inheritance as Weismannian inheritance can, a larger diversity in X must also be maintained.

**Figure SI 11.**
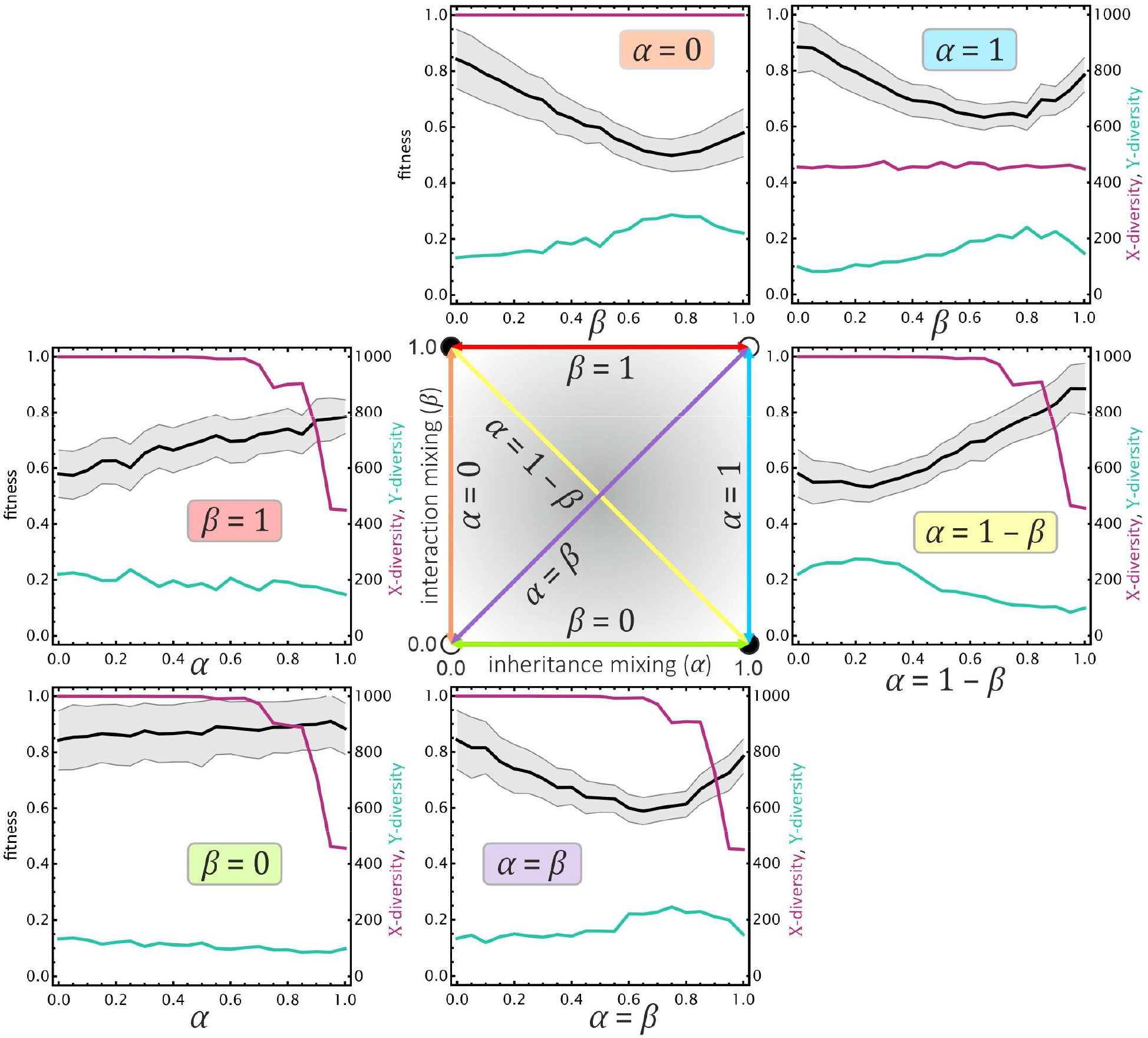
Parameter scans across the Lamarckian-Weismannian continuum of Figure 6. The continuum is represented by the square in the middle, the coloured arrows correspond to plots around it with the corresponding coloured labels. Plots show the average fitness of the population (left axis) and the X and Y sequence diversities (number distinct sequences) of the population (right axis). Each plot point is the average of the equilibrium of the simulation (i.e., average of 50 equidistant points sampled from the last 10 000 replication steps per simulation). Parameters are: N_X_ = 30, N_Y_ = 10, c_X_ = 3, c_Y_ = 1, a_X_ = a_Y_ = 2, θ_XY_ = θ_uniformly ordered_, 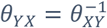, μ_XX_ = μ_XY_ = μ_YX_ = 0.01, population size of 1000, and 100 000 replication steps.

### 7. Language learning as an example of cultural inheritance

Let us demonstrate our argument that cultural learning is practically informational replication through the example of language learning according to ^34^ (see Figure SI 12). Suppose a human language user has a *hypothesis h* of a language, represented as a set of rules in her brain (e.g. constructions ^35,37,141^). Her “total” language *L* is the set of all possible utterances that can be generated from this set of rules. To learn the language, the learner must infer the rules. Grammar is not learnt by copying explicit grammatical rules directly (especially not by children and presumably not by early *Homo*) but through a limited number of incomplete or partial linguistic examples ^142^ (cf. poverty of stimulus ^34^). Presumably, if a learner would be able to process all *L*, he could, in theory, reconstruct all *h*. Alas, *L* is expected to be astronomical in size and no speaker ever is expected to produce all of it. Real language learners make do with much less. Any learner only receives a small subset of the *L*, the realized language (and even that may be corrupted). A learner attempts to reconstruct *L* by *inferring* the original rules as *h*^*^. Due to the lack of template copying and the information-losing nature of utterance generation, it is likely that *h*^*^ ≠ *h*, even when assuming zero errors during production and learning. It seems then, that there can be no exact replication of the internal language *h*.

**Figure SI 12.**
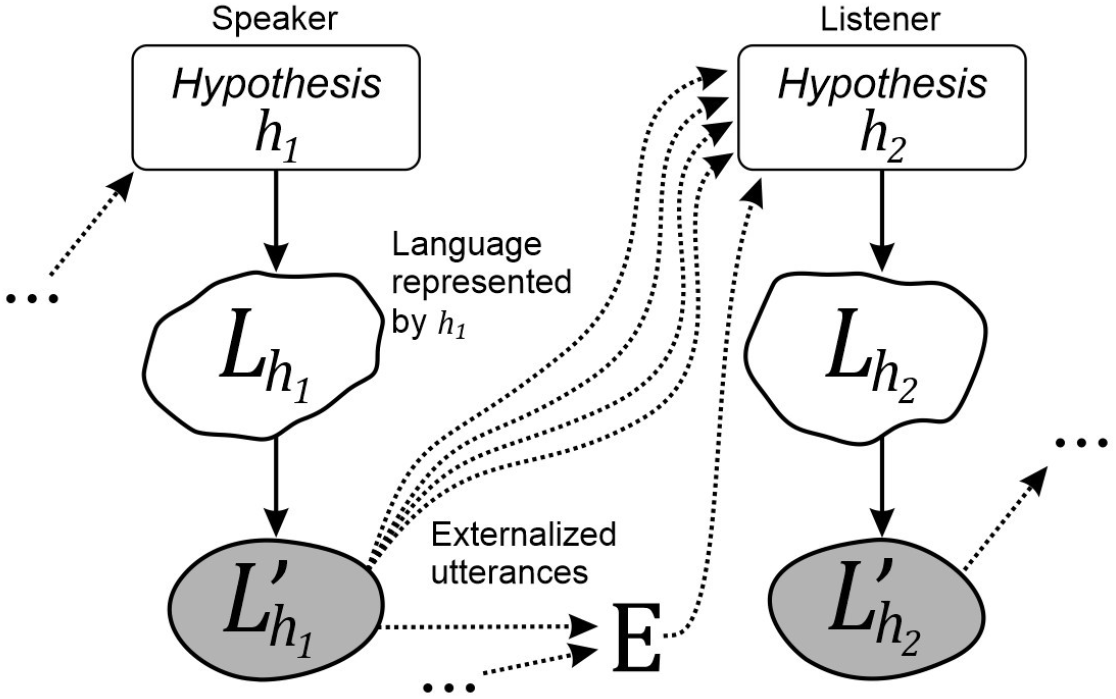
A model of language learning adapted to the topology of cultural inheritance of Figure 2C (the iterated learning model, reproduced from ^34^). Suppose a human language user, Agent 1, has a hypothesis of the language, represented as a set of rules, h_l_ = {x_l_, x_2_, … }. Agent 1’s “total” language is the set of all possible utterances L = {y_l_, y_2_, … } that can be generated from h. Due to the limited and constrained nature of real languages, the learner receives only a small set of utterances L′ ⊂ L (dashed arrows) as examples of L, which is the realized language. He then reconstructs L by inferring the original ruleset h as h^*^. The lack of direct template copying gives room for informational decay, and it is likely that h^*^ ≠ h. This might jeopardize the mutual understanding of language users. However, the iterative nature of language learning involves not only multiple, repeated examples (multiple dashed arrows), but potentially external sources E generated by the speaker or other language users (e.g. utterances, books, blueprints, etc.) that can further help the learner to successfully reconstruct L by inference. Ultimately, h^*^ will be phenotypically identical to h, which ensures that 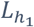 and 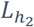 are sufficiently similar for linguistic continuity among users.

However, as we have established above, the successive external states *h* and *h*^*^ need not be identical for language users to understand each other. As long as they both encode the same external language, or more strictly speaking, *h*^*^ is phenotypically equivalent to *h* (*h*∼*h*^*^), realized languages are equivalent 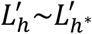. That is, under reasonable selection criteria, linguistic continuity is maintained. In plain words, if the two different grammar sets in two individuals generate languages with which they can understand each other, then language is maintained and *L*′ is replicated from time to time (ignoring mutations for sake of simplicity). Crucially, due to the Lamarckian information transmission involved, the two internal representations *h* and *h*^*^ may differ substantially, and they are only equivalent because *they generate functionally identical languages*. We expect that internal concepts (e.g., lexicon, grammar, colours, tools, etc.) are represented differently in different brains despite being phenomenologically (i.e., externally) equivalent.

### 8. Supplementary references

TBA

